# An ULK1/2-PXN mechanotransduction complex suppresses breast cancer cell migration

**DOI:** 10.1101/2023.02.03.526950

**Authors:** Peigang Liang, Jiaqi Zhang, Yuchen Wu, Shanyuan Zheng, Zhaopeng Xu, Shuo Yang, Jinfang Wang, Suibin Ma, Li Xiao, Tianhui Hu, Wenxue Jiang, Qiong Xing, Mondira Kundu, Bo Wang

## Abstract

The remodeling and stiffening of the extracellular matrix (ECM) associated with breast cancers is a well-recognized modulator of disease progression. However, how changes in the mechanical properties of the ECM are converted into biochemical signals that direct tumor cell migration and metastasis remains poorly characterized. Here, we describe a new role for the autophagy-inducing serine/threonine kinases ULK1 and ULK2 in mechanotransduction. We demonstrate that ULK1/2 activity inhibits the assembly of actin stress fibers and focal adhesions (FAs), and as a consequence impedes cell contraction and migration. Mechanistically, we identify PXN/paxillin, a key component of the mechanotransducing machinery, as a direct binding partner and substrate of ULK1/2. ULK-mediated phosphorylation of PXN at S32 and S119 weakens homotypic interactions and liquid-liquid phase separation of PXN, impairing FA assembly, which in turn impedes the mechanotransduction of breast cancer cells. ULK1/2 and the well characterized PXN regulator, FAK/Src, have opposing functions on mechanotransduction and compete for phosphorylation of adjacent serine and tyrosine residues. Thus, our study reveals ULK1/2 as important regulators of PXN-dependent mechanotransduction.

**Highlights:**

- ULK1/2 interact with PXN and phosphorylate PXN at S32 and S119 in response to mechanical stimuli
- ULK1/2-mediated phosphorylation of PXN regulates mechanotransduction and migration of breast cancer cells
- ULK1/2 modulate the biomaterial properties of focal adhesions through PXN phosphorylation
- ULK1/2 and FAK/Src act antagonistically in mechanotransduction through competitive phosphorylation of PXN

## Introduction

The extracellular matrix (ECM) continuously modulates various aspects of cellular behavior, including division, growth, migration, and death. Cells sense changes in the biochemical composition or mechanical properties of the ECM through focal adhesions (FAs), which are dynamic protein assemblies involved in mechanotransduction. Upon engagement with the ECM, transmembrane heterodimeric integrin receptors recruit numerous FA-associated components, which in turn connect with the actin cytoskeletal network. In response to increasing stiffness, cells actively strengthen opposing contractile and tension forces through actin polymerization and actomyosin motors. In this manner, mechanical cues are converted into biochemical outputs, which include extensive post-translational modifications and changes in protein-protein interactions.

Tumor-associated breast tissues exhibit augmented rigidity compared to normal mammary tissues. Indeed, palpation is used as an initial diagnostic method for breast tumors due to their prominently rigid nature. It is now widely appreciated that stiffened ECM instructs invasion of breast cancer cells (Levental et al., 2009; Riehl et al., 2020). However, the molecular mechanisms through which breast cancer cells sense and transduce these mechanical signals into biological activities remain less well understood.

Most steps in mechanotransduction pathways are energy consuming, requiring approximately 50% of total cellular ATP (Bernstein and Bamburg, 2003). Therefore, these pathways are likely to engage in crosstalk with metabolic programs (Romani et al., 2021; Salvi and DeMali, 2018). In principle, metabolism can impinge on mechanotransduction in two distinct manners. First, energy-producing catabolism can fuel processes such as actin polymerization. Alternatively, certain metabolites can directly modulate cellular processes that are essential for mechanotransduction. Preliminary evidence suggests that mechanotransduction and metabolism can be coupled. For instance, the metabolic sensor AMPK is activated in response to mechanical stimuli and stimulates both ATP production and glucose transport (Bays et al., 2017), which may in turn reinforce cell adhesions and the actin cytoskeletal network. However, whether AMPK-mediated glucose transport and ATP production or other targets of AMPK, such as ULK1/2, are responsible for the enhanced cell-cell contact remains unclear.

ULK1/2 are functionally redundant serine/threonine kinases that regulate multiple steps of the autophagy pathway, which targets superfluous or damaged proteins and organelles for lysosomal degradation and recycling to maintain cellular energy homeostasis (Dikic and Elazar, 2018; Wang and Kundu, 2017; Zachari and Ganley, 2017). Phosphorylation of ULK1/2 by AMPK increases flux through the autophagy pathway (Kim et al., 2011). The inhibition of ULK1-dependent autophagic function under hypoxic conditions has been associated with increased invasion and migration of breast cancer cell lines *in vitro* and in xenograft models (Dower et al., 2017), and although autophagy has been shown to promote FA disassembly and cell motility (Bressan et al., 2020; Kenific et al., 2016; Sharifi et al., 2016), the role of ULK1/2 in mechanotransduction has not been explored. Indeed, the links between mechanotransduction and autophagy remain controversial. Whereas autophagy activity is enhanced via nuclear-localized YAP/TAZ induced by high ECM stiffness (Totaro et al., 2019), autophagy can also be increased in cells grown on soft matrix (Vera-Ramirez et al., 2018). These observations highlight a context-dependent relationship between these two pathways. Not surprisingly, autophagy can be anti- or pro-tumorigenic depending on the tumor stage and context (Niklaus et al., 2021).

In examining the relationship between autophagy and mechanotransduction, we made the unexpected discovery that ULK1 and ULK2 negatively regulate mechanotransduction in breast cancer cells. Specifically, we demonstrate that ULK1/2 interact directly with the FA scaffold protein PXN/paxillin, and phosphorylate PXN at S32 and S119 in a mechanosensitive manner. Phosphorylation of PXN by ULK1 weakens homotypic interactions, thereby reducing PXN’s ability to drive the assembly of FA condensates *in vitro*, and decelerating the kinetics of FA assembly in cells, thus compromising mechanotransduction and migration. These observations may help to explain why decreased expression of ULK1 in breast cancer samples is associated with a worse prognosis.

## Results

### ULK1/2 negatively regulate mechanics of breast cancer cells in an autophagy-independent manner

In order to begin exploring the relationship between the ECM, mechanotransduction and autophagy, we collected 6 samples from breast cancer patients along with paired adjacent normal tissues and performed immunoblot analyses. Consistent with previous report (Levental et al., 2009), we observed increased levels of certain ECM proteins, including collagen and fibronectin, in the tumor samples compared to normal tissues (Fig. 1A–1B). Consistent with the results of our sc-RNAseq analyses, tumor samples also showed increased levels of several markers of cellular contraction and integrin signaling, such as α-smooth muscle actin (α-SMA), focal adhesion kinase (FAK) phosphorylated at Y397, and PXN phosphorylated at Y118 (Fig. 1A–1B). We found no detectable change in the protein levels of several key markers (ATG13, FIP200/RB1CC1, p62, and LC3II) of autophagic activity, except ULK1, whose levels were decreased (Fig. S1A–1B).

**Figure 1.**
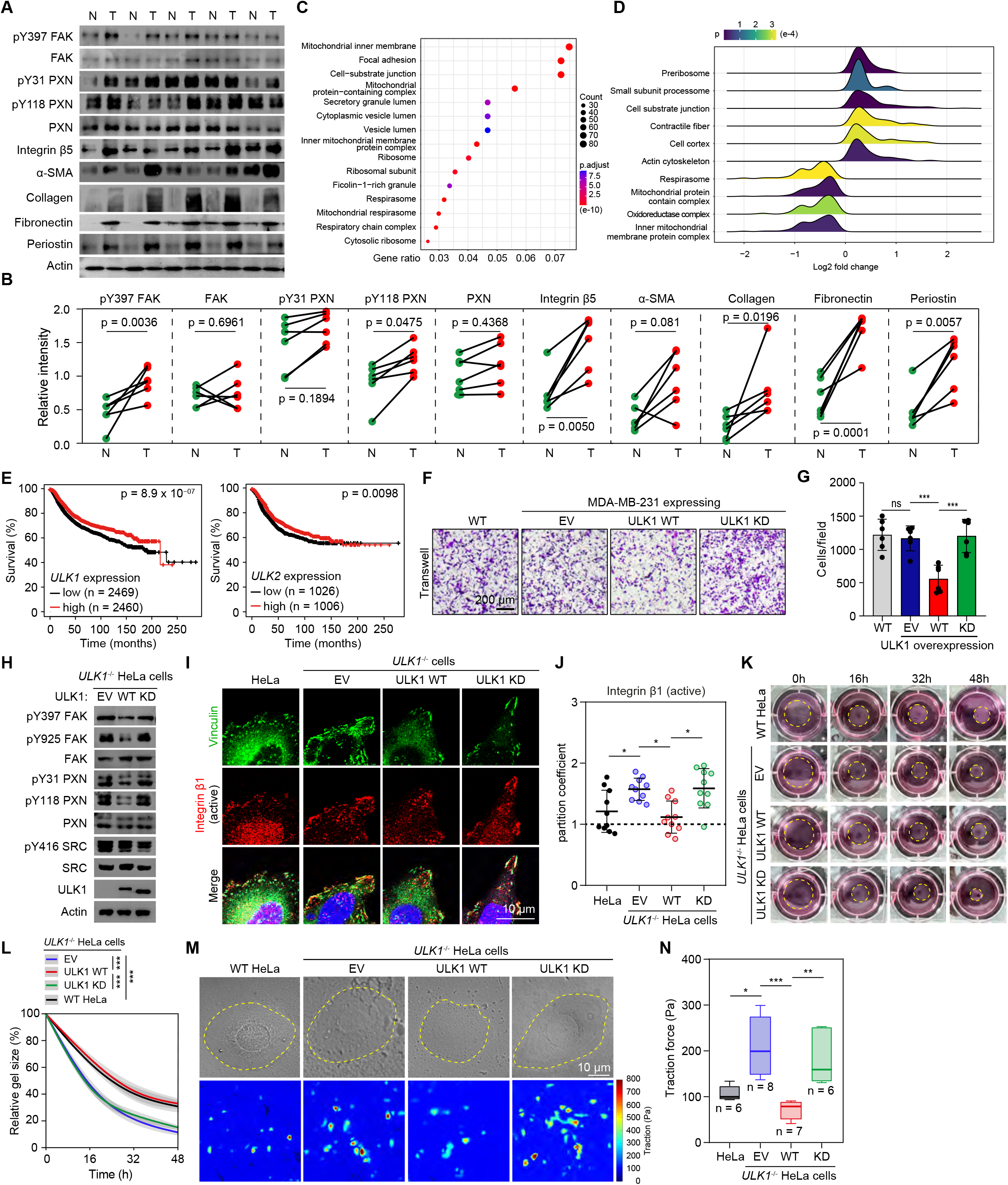
ULK1/2 regulate cell migration and mechanotransduction of breast cancer. (**A**, **B**) Lysates prepared from normal (N) and breast tumor (T) tissues were analyzed by immunoblot with the indicated antibodies. The relative band intensities were quantified by densitometry (**B**). Data are presented as mean ± SEM from 6 individual patients. Statistic difference was calculated by paired Student’s t-test. (**C**) The 15 most significantly enriched pathways yielded from Gene Ontology analyses from deposited sc-RNAseq datasets of both normal and breast cancer patients. (**D**) Gene Set Enrichment Analysis of the same sc-RNAseq datasets. The 10 most significantly enriched pathways were shown. (**E**) The correlation of ULK1/2 RNA expression levels and survival of breast cancer patients. (**F**) Migratory potentials of WT or *ULKT*^-/-^ MDA-MB-231 cells were assessed by transwell assay. The cells migrated to the lower chamber were stained with crystal violet (**F**). Quantification of the cell number was show in (**G**). Data are presented as mean ± SEM. (**H**) Cell lysates prepared from *ULK1*^-/-^ HeLa cells expressing EV, ULK1 WT or KD were analyzed by immunoblot using the indicated antibodies. (**I**, **J**) WT or *ULKT*^-/-^ HeLa cells reconstituted with EV, ULK1 WT or KD were immunostained with antibodies against active integrin β1 (12G10) and Vinculin. Representative confocal images were shown in (**I**), and the relative intensities of active integrin β1 in the FAs were quantified in (**J**). (**K**, **L**) WT or *ULK1*^-/-^ HeLa cells reconstituted with EV, ULK1 WT or KD were seeded in 3D collagen gels. The gel size at each time point was photographed (**K**) and measured (**L**). Data are shown as mean ± SEM from 3 independent experiments. (**M**, **N**) WT or *ULK1*^-/-^ HeLa cells reconstituted with EV, ULK1 WT or KD were analyzed by TFM. The total cellular force was quantified in (**N**). ns, not significant; *p < 0.05; **p < 0.01; ***p < 0.001 by one-way ANOVA.

Due to the well-known cellular complexity of tissues infiltrated by breast cancer, and heterogeneity of breast cancer cells, we sought to identify consistent cell intrinsic differences between normal and malignant breast epithelial cells. Towards this end, we used a publicly available single-cell RNA sequencing (sc-RNAseq) dataset to compare gene expression profiles from normal and malignant mammary epithelial cells after computationally separating them from other cell types in the heterogeneous tissue samples (Pal et al., 2021). Gene ontology (GO) enrichment revealed 111 significantly altered cellular processes. Among the top 15 most significant hits, we found prominent enrichment of mitochondria and ribosome-related activities (Fig. 1C), both of which are frequently associated with breast cancers (Ebright et al., 2020; Li and Li, 2021). Interestingly, the top GO terms also included FA and cell substrate junction, both of which are directly relevant to cellular mechanosensing and transduction (Fig. 1C). We next performed gene set enrichment analysis (GSEA), which revealed that genes related to cell substrate junction, contractile fiber, cell cortex, and actin cytoskeleton were significantly upregulated in the malignant mammary epithelial cells (Fig. 1D). These data highlight the aberrant activation of the mechanotransduction pathway in breast cancer cells compared to normal mammary epithelium. Notably, the abundance of important autophagy-associated genes remained unchanged between normal and malignant cells (Fig. S1C).

The reduced levels of ULK1 in cancerous breast tissues drew our attention because ULK1 expression has been reported as an independent prognostic factor in breast cancer (Tang et al., 2012). Using the TCGA database, we independently confirmed the association between the expression of ULK1 and ULK2 and breast cancer prognosis [i.e., decreased *ULK1* and *ULK2* mRNA levels were associated with a significantly (p = 8.9 x 10^-07^ and 0.0098 for *ULK1* and *ULK2*, respectively) worse prognosis] (Fig. 1E).

To test the effect on ULK1 expression on cell migration, we used a chemoattractant-based transwell assay in which cells are cultured in the upper compartment and allowed to migrate through a microporous membrane into the lower compartment, where chemotactic agents are present. Since highly metastatic breast cancer cells such as MDA-MB-231 exhibit relatively low levels of ULK1 expression compared to those that are less metastatic (Mao et al., 2020), we chose to inducibly express ULK1 wildtype (WT), ULK1 kinase dead (KD, K46A), ULK2 WT, or ULK2 KD (K39T) in these cells. The steady state levels of ULK1 (WT and KD) were higher than that of ULK2 (WT or KD) in the stable cell lines (Fig. S1D). Overexpression of ULK1 WT, but not ULK1 KD, strongly impeded cell migration in this assay (Fig. 1F–1G) without any appreciable impact on cell proliferation (Fig. S1E). We also generated *ULK1* null HeLa cells, reconstituted them with ULK1 WT or ULK1 KD (Fig. S1F), and assessed the migratory potentials of these cells using the transwell assay. Again, ULK1 WT, but not ULK1 KD, inhibited cell migration in these cells (Fig. S1G–1H), suggesting that this function of ULK1 in tumor cell migration is likely not cell type-specific. Silencing either *ULK1* or *ULK2* markedly increased migration of HeLa cells (Fig. S1I–1J), highlighting their functional redundancy.

To test whether this inhibitory effect of ULK1/2 in cell migration is related to its effects on autophagy, we chose to knock down expression of several key autophagy genes, including *ATG13*, *FIP200/RB1CC1*, *ATG7*, and *ATG14*, and assess the subsequent impact on cell migration (Fig. S1K). Among these, ATG13 and FIP200 form complexes with ULK1/2, ATG7 is required for most known forms of autophagy, and ATG14 is a substrate of ULK1/2 during autophagy initiation (Dikic and Elazar, 2018; Park et al., 2018). Knockdown of *ATG13*, *ATG7*, or *ATG14* did not significantly alter cell migration, whereas knockdown of *FIP200* significantly increased cell migration (Fig. S1L–1M). Although recent studies have focused on FIP200’s role in autophagy, FIP200 was initially identified as a direct interactor and regulator of the FA components FAK and Pyk2 (Ueda et al., 2000). Indeed, FIP200’s regulation of cell motility and focal adhesion stability is independent of its role in autophagy (Abbi et al., 2002; Assar and Tumbarello, 2020). Taken together, these results suggest that ULK1/2 negatively regulate breast cancer cell migration in an autophagy-independent manner.

Because MDA-MB-231 cells are known for extreme morphological and mechanical heterogeneity (Shen et al., 2020), for subsequent studies we used HeLa cells for morphological analyses and performed critical functional assays (e.g., transwell) using MDA-MB-231 cells to test the applicability of our findings in breast cancer cells.

Given our finding of the negative correlation between integrin signaling and ULK1 protein levels in patient cells, we hypothesized that ULK1’s inhibition of cell migration may at least in part be related to an inhibitory effect on integrin activation. To test this concept, we next examined signaling proteins downstream of integrin activation in response to manipulation of ULK1 expression or activity. Indeed, overexpression of ULK1 WT was associated with decreased abundance of signaling proteins downstream of integrin activation, including FAK phosphorylated at Y397 and Y925 as well as PXN at Y31 and Y118 (Fig. 1H, and S1N). We next asked if the overall activity of integrin receptors was altered by ULK1. We took advantage of a well-characterized antibody (12G10) that specifically recognizes the activated integrin β1 receptor. Indeed, depleting ULK1 led to significantly increased intensities of activated integrin β1 receptor and re-introducing the kinase-active ULK1 attenuated the intensities (Fig. 1I–1J). Because integrin signaling is tightly associated with cell contraction, we performed a 3D collagen contraction assay, which measures the cellular contractile forces towards the matrix, to assess the role of ULK1 in this process. Depleting ULK1 significantly diminished cellular contractile forces, which was reversed by reconstituting kinase active ULK1 (Fig. 1K–1L). Again, this function of ULK1 was unrelated to autophagy since knockdown of other representative autophagy proteins did not have similar effects (S1O-1P). We next employed quantitative traction force microscopy (TFM) to evaluate the average cellular contractile force, which was reduced from 200 Pa in *ULK1*^-/-^ cells to less than 100 Pa by the expression of functional ULK1 (Fig. 1M–1N). Collectively, these results suggest that ULK1 is a negative regulator of cell contraction and migration.

### ULK1/2 suppress stiffness-dependent cell spreading, actin reinforcement and FA assembly

Based on the findings described above, we hypothesize that ULK1/2 govern cellular response to mechanical changes of the ECM. To mimic the fluctuating mechanical stimulation of the ECM, we fabricated polyacrylamide (PA) gels with various stiffness ranging from 2.7 kPa (soft) to glass (stiff) and used these as substrates for HeLa cell growth. We then collected lysates from these cells and examined markers of cell contraction and integrin signaling by immunoblot. As expected, we found upregulation of α-SMA, phosphorylated myosin light chain 2 (MLC2), FAK, and PXN with increasing stiffness of the substrate (Fig. 2A–2B). We also examined metabolic proteins to investigate potential association between mechanotransduction and metabolism. Among proteins we examined, AMPK was significantly activated with increasing substrate stiffness (Fig. 2A–2B), consistent with the idea that AMPK safeguards the energetic needs for resisting increases in substrate stiffness (Bays et al., 2017). mTOR activity was also markedly augmented by higher substrate stiffness, in line with the idea high substrate stiffness promotes cell growth (Fig. 2A–2B). In contrast to prior studies showing changes in autophagic activity in response to increases in substrate stiffness (Totaro et al., 2019; Vera-Ramirez et al., 2018), we found that autophagy flux remained stable, as indicated by unchanged levels of p62, LC3II, and phosphorylated ATG14 protein (Fig. 2A–2B). Morphologically, cells expressing ULK1 WT showed significantly shrunken size and disrupted actin remodeling compared to those expressing no ULK1 or KD ULK1 (Fig. 2C-E, and S2A), suggesting that ULK1 negatively governs cellular mechanics in a kinase-dependent manner. Knockdown of *ATG13*, *FIP200/RB1CC1*, *ATG7*, or *ATG14* had no appreciable effects on cell spreading and actin structure, whereas knockdown of *ULK1* resulted in over-spreading and actin super-assembly (Fig. S2B-D), implying that ULK1 participate in mechanotransduction pathways independent of autophagy. We found similar effects of ULK2 (i.e., *ULK2*^-/-^ cells with re-expression of ULK2 WT and KD) in regulating cell size and actin structure (Fig. 2F–2G, and S2E). Live cell imaging revealed that ULK1-expressing cells showed delays in cell spreading on glass (Fig. S2F–2G). This phenomenon was independently validated by silencing ULK1 and ULK2 using shRNAs (Fig. S2H). For cells growing on glass, the decreased actomyosin contraction due to ULK1 expression was accompanied by lower cellular stiffness measured by atomic force microscopy (AFM) (Fig. 2H). Together, our data implicate an intriguing role of ULK1/2 in negatively regulating actin assembly, leading to reductions in cell stiffness, contractile forces and adaptability to mechanical inputs.

**Figure 2.**
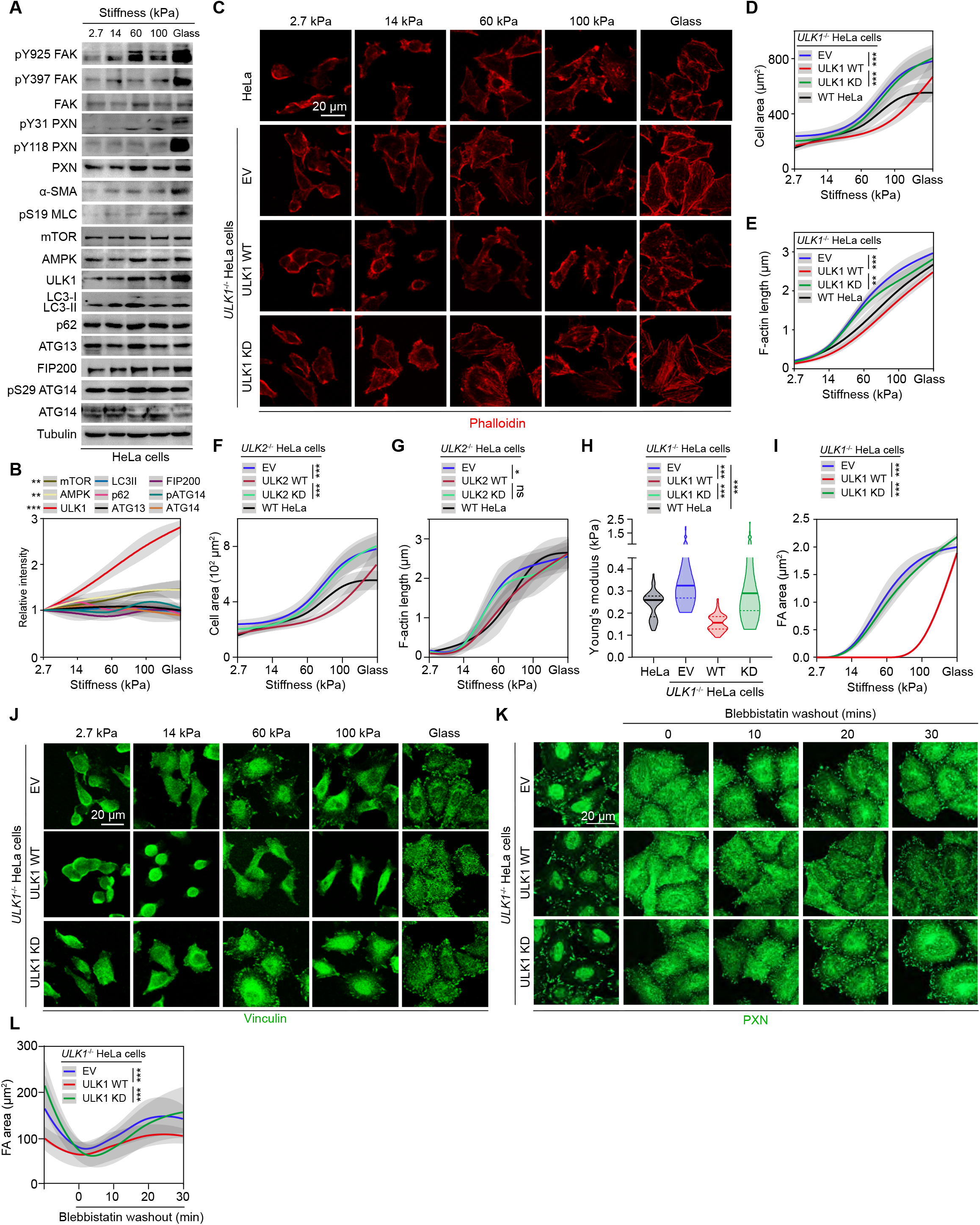
ULK1/2 suppress stiffness-dependent cell spreading, actin reinforcement and FA assembly. (**A**, **B**) Lysates prepared from WT HeLa cells grown on surfaces with different stiffness were analyzed by immunoblot with the indicated antibodies (**A**). The relative band intensities were quantified by densitometry (**B**). Data are presented as mean ± SEM from 3 independent experiments. (**C**-**E**) WT or *ULKT*^-/-^ HeLa cells expressing EV, ULK1 WT or KD were grown on surfaces with different stiffness. The cells were stained with Phalloidin (**C**). The cell area (median ± 95% confidence interval) and F-actin length (mean ± SD) were quantified and shown in (**D**) and (**E**), respectively. (**F**, **G**) WT or *ULK2*^-/-^ HeLa cells expressing EV, ULK2 WT or KD were grown on surfaces with different stiffness. The cell area (median ± 95% confidence interval) and F-actin length (mean ± SD) were quantified and shown in (**F**) and (**G**), respectively. (**H**) Young’s modulus of WT or *ULK1”* HeLa cells expressing EV, ULK1 WT or KD was measured by AFM. (**I**, **J**) *ULK1*^-/-^ HeLa cells expressing EV, ULK1 WT or KD cultured on surfaces with different stiffness were immunostained with Vinculin antibody to visualize FAs (**J**). The size of FAs under each individual condition was quantified (**I**). Data are presented as median ± 95% confidence interval. (**K**, **L**) *ULKT*^-/-^ HeLa cells expressing EV, ULK1 WT or KD were treated with DMSO or blebbistatin. The cells were fixed at different time points after Cytochalasin D washout, and FAs were visualized by PXN immunostaining (**K**). Total FA area was quantified. Data are presented as median ± 95% confidence interval. ns, not significant; *p < 0.05; **p < 0.01; ***p < 0.001 by one-way ANOVA.

FAs are dynamic transmembrane macromolecular assemblies serving as a major node to translate mechanical stimuli to biological signal and dictate downstream cellular behavior, including cell migration, differentiation, and division (Parsons et al., 2010). Based on our findings that ULK1/2 negatively regulate cell mechanotransduction, we next tested the hypothesis that ULK1/2 influence the dynamics of FAs. Consistent with this hypothesis, stiffer substrates significantly increased the formation of FAs, which was markedly attenuated by expression of ULK1 WT but not ULK1 KD (Fig. 2I–2J). We next treated cells expressing different ULK variants grown on glass with the myosin inhibitor blebbistatin and allowed the cells to recover while we monitored the assembly/maturation of FAs. We found that ULK1 significantly hindered FA assembly/maturation (Fig. 2K–2L). In contrast, knockdown of *ATG13*, *FIP200/RB1CC1*, *ATG7*, or *ATG14* resulted in relatively modest changes in FA morphology as compared to *ULK1* knockdown (Fig. S2I–S2K). Based on these results, we propose that ULK1/2 suppress stiffness-dependent cell spreading, actin reinforcement, and FA assembly via autophagy-independent mechanisms.

### ULK1/2 interact with and phosphorylate PXN directly both in vitro and in cells

To pursue the molecular mechanism by which ULK1/2 has these effects, we first examined PXN, a well-studied adaptor protein within FAs that has been shown to genetically interact with ULK1/2 in *Drosophila* (Chen et al., 2008). PXN is a highly conserved protein that harbors 5 LD domains in its N terminus and 4 LIM domains in its C terminus. Both the N and C termini mediate protein-protein interactions that are implicated in FA dynamics and mechanotransduction (Deakin and Turner, 2008).

We began by testing interactions between ULK1/2 and PXN by immunoprecipitation in HEK293T cells. We found that exogenously expressed PXN co-immunoprecipitated with ULK1/2 (Fig. 3A). Recombinant PXN also pulled down with ULK1/2 immunopurified from HEK293T cells, suggesting direct interactions (Fig. 3B). We next generated a series of tagged ULK1 and PXN truncation mutants and expressed these proteins in HEK293T cells (Fig. 3C). We found that ULK1/2 co-immunoprecipitated with the LIM4 domain located within the C terminus of PXN (Fig. 3D–3E, and Fig. S3A), and that full-length PXN co-immunoprecipitated with the C terminus of ULK1 (Fig. 3F). These results prompted us to examine whether ULK1/2 were also present in FAs along with PXN. Indeed, both ULK1 and ULK2 colocalized with PXN in the FAs in HeLa cells (Fig. 3G–3H).

**Figure 3.**
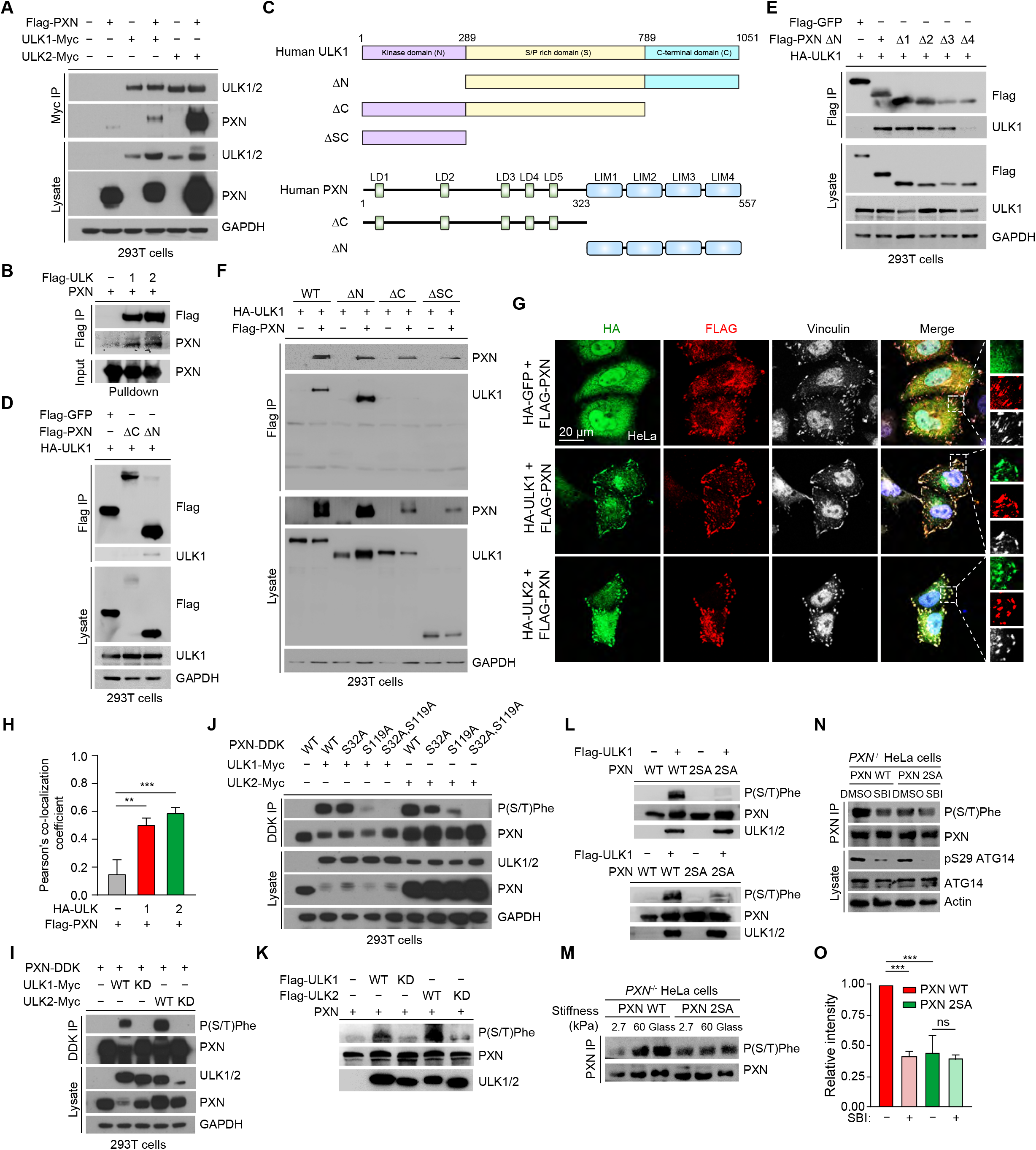
ULK1/2 directly interact with and phosphorylate PXN at S32 and S119, in response to mechanical stimuli. (**A**) 293T cells were transfected with the indicated constructs and subjected to Myc immunoprecipitation. (**B**) Pull-down assays using ULK1 and ULK2 immunopurified from 293T cells and recombinant PXN. (**C**) Schematic illustrations of the different constructs of both ULK1 and PXN. (**D**-**F**) 293T cells transfected with the indicated constructs were subjected to Flag immunoprecipitation. (**G**) WT HeLa cells were transiently co-transfected with Flag-PXN and HA-GFP, HA-ULK1, or HA-ULK2. Cells were fixed and processed for immunostaining using Flag and HA antibodies. (**H**) Quantification of the colocalization between PXN and ULK1/2 from (**G**). Data are presented as mean ± SD. (**I**) Lysates prepared from 293T cells transfected with the indicated constructs were analyzed by immunoblot with the indicated antibodies. (**J**) Lysates prepared from 293T cells transfected with different PXN mutants together with ULK1 or ULK2 were analyzed with the indicated antibodies. (**K**) *In vitro* kinase assay was performed using ULK1 or ULK2 immunopurified from 293T cells and recombinant PXN. (**L**) *In vitro* kinase assay using ULK1 or ULK2 and PXN WT or 2SA mutant. (**M**) *PXN*^-/-^ HeLa cells reconstituted with PXN WT or 2SA were seeded on surfaces with different stiffness. Immunoprecipitated PXN was analyzed by immunoblot with P(S/T)Phe antibody. (**N**) *PXN*^-/-^ HeLa cells reconstituted with PXN WT or 2SA growing on glass were treated with DMSO or SBI-0206965 (SBI). Immunoprecipitated PXN was analyzed by immunoblot with P(S/T) antibody. (**O**) The relative band intensities of P(S/T)Phe from (**N**) were quantified by densitometry. Data are presented as mean ± SEM, n = 3 independent experiments. ns, not significant; **p < 0.01; ***p < 0.001 by one-way ANOVA.

We also examined interactions between ULK1/2 and HIC5, a close paralogue of PXN sharing extensive sequence and structural similarities (Alpha et al., 2020; Deakin and Turner, 2008). Both redundant and non-redundant functions have been reported for PXN and HIC5 (Alpha et al., 2020; Deakin and Turner, 2008). Not surprisingly, HIC5 also interacted with ULK1 and ULK2 in a fashion akin to PXN (Fig. S3B–S3C), and C-terminal regions of HIC5 and ULK1 were responsible for these interactions (Fig. S3D–S3E). Given the homology and functional redundancy between PXN and HIC5, we focused our subsequent studies on PXN.

Proteins interacting with ULK1/2 are often direct substrates of these kinases (Wang and Kundu, 2017). We took advantage of a previously described p(S/T)Phe antibody that recognizes phosphorylated serine/threonine residues with tyrosine, tryptophan, or phenylalanine at the −1 position or phenylalanine at the +1 position. When phosphorylated, the target serine/threonine sites of many substrates of ULK1/2 can be detected by this antibody (Joo et al., 2016; Russell et al., 2013). We transiently co-expressed PXN with ULK1, ULK2, and their KD mutants in HEK293T cells and found that PXN was phosphorylated by both ULK1 and ULK2 in a kinase-dependent manner (Fig. 3I). Human PXN harbors 8 serine/threonine sites that when phosphorylated, can be detected by the p(S/T)Phe antibody. To determine which of these sites were targets of ULK1/2 phosphorylation, we performed site-directed mutagenesis to substitute each of these residues with alanines (S32A, S89/90/91A, S119A, S164A, S382A, T475A, S481A, T540A). We found that that alanine substitutions at S32 and S119 dramatically diminished ULK1-mediated phosphorylation of PXN in HEK293T cells (Fig. S3F). As expected, a 2SA (S32A, S119A) PXN mutation was sufficient to abolish the phosphorylation (Fig. 3J). We next performed *in vitro* kinase assays, finding that ULK1/2-mediated phosphorylation of PXN at S32 and S119 was direct (Fig. 3K–3L). We attempted to generate antibodies raised against phospho-peptides containing S32 and S119, respectively, but did not succeed (data not shown). Therefore, we continued to utilize the p(S/T)Phe antibody to characterize PXN phosphorylation by ULK1/2. Mechanical stimulation with increasing substrate stiffness elicited striking upregulation of PXN phosphorylation (Fig. 3M), which was attenuated by a selective ULK inhibitor, SBI-0206965 (Fig. 3N–3O). Furthermore, the PXN 2SA mutant was refractory to mechanical stimuli-induced phosphorylation (Fig. 3M–3O). When cells growing on still glass were treated with myosin inhibitor blebbistatin, PXN phosphorylation was dramatically reduced (S3G-S3H). Conversely, when cells growing on relatively soft substrate (60 kPa) were stimulated with RhoA activator (Rho activator II), PXN phosphorylation was markedly enhanced. Collectively, our data demonstrate that ULK1/2 directly phosphorylate PXN at S32 and S119 *in vitro* and likely in cellular mechanotransduction as well.

### ULK1/2-mediated phosphorylation of PXN gate-keeps cellular mechanotransduction

To determine the functional impact of ULK-mediated phosphorylation of PXN, we stably expressed EV (empty vector), PXN WT, the phospho-defective 2SA, or a phospho-mimetic 2SD and assessed their ability to restore the aberrant migration and mechanotransduction of PXN null cells. In MDA-MB-231 cells, PXN WT efficiently rescued the compromised migration of *PXN*^-/-^ cells, whereas expression of PXN 2SD failed to do so (Fig. 4A–4B). Such inhibitory effects of PXN phosphorylation on cell migration were similarly observed in HeLa cells (Fig. S4A–S4B). More importantly, the PXN 2SD mutant was capable of restraining cell migration even in cells depleted of ULK1 by shRNA (Fig. S4C–S4D), suggesting that PXN functions downstream of ULK1 in this pathway.

**Figure 4.**
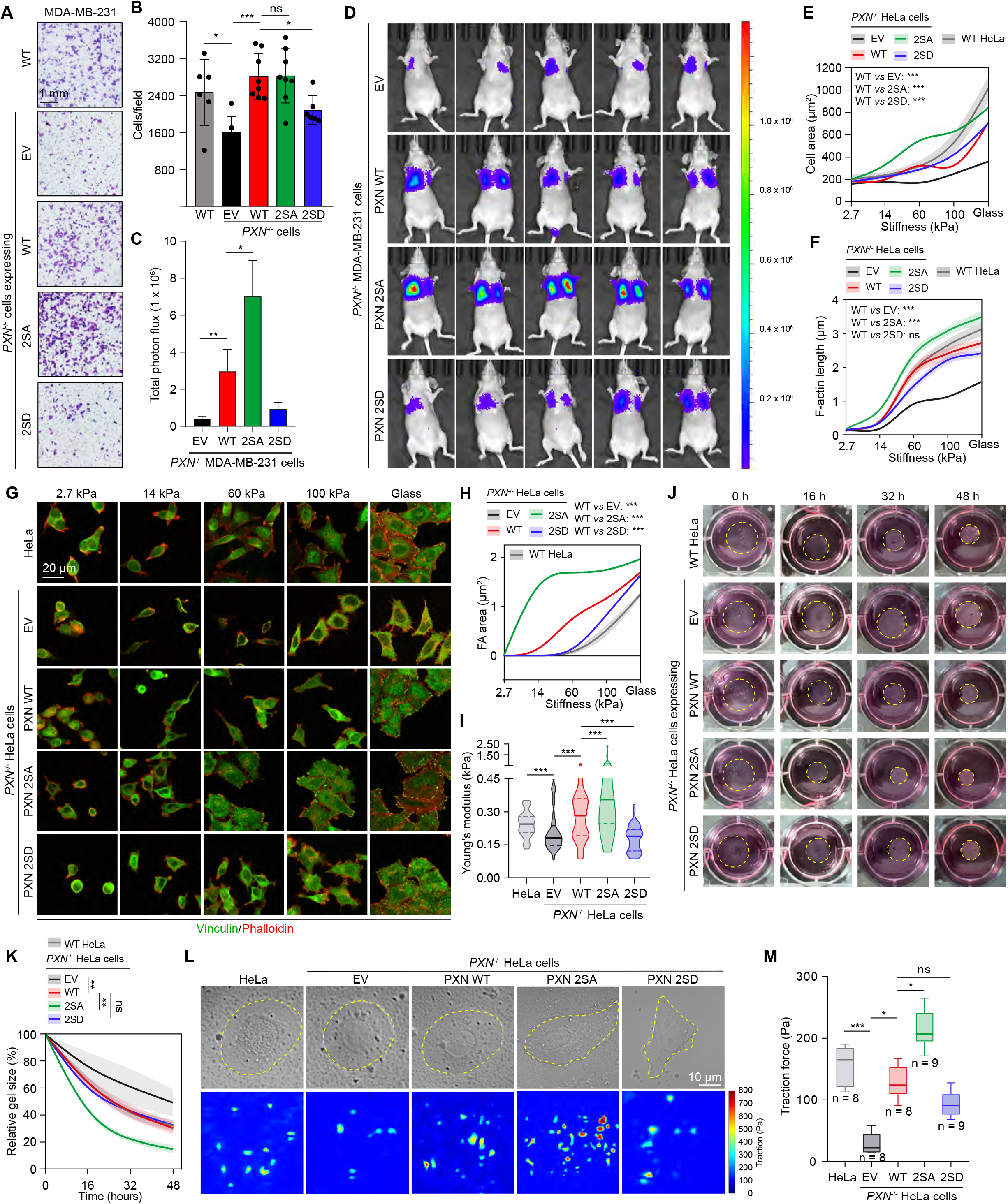
ULK1/2-mediated phosphorylation of PXN inhibits breast cancer cell mechanics. (**A**, **B**) WT or *PXN*^-/-^ MDA-MB-231 cells reconstituted with EV, PXN WT, 2SA, or 2SD were subjected to transwell assay. The cells were stained with crystal violet (**A**). Quantification of cells migrated to the lower chambers was show in (**B**). Data are presented as mean ± SEM. (**C**, **D**) *PXN*^-/-^ MDA-MB-231 cells reconstituted with EV, PXN WT, 2SA, or 2SD were injected into nude mice through tail vein. Cells migrating to the lung were imaged with luciferase. Quantification of total photo flux per mouse were shown in (**C**). n = 7, 6, 7, and 5 for EV, PXN WT, 2SA, and 2SD, respectively. (**E-H**) WT or *PXN*^-/-^ HeLa cells expressing EV, PXN WT, 2SA, or 2SD were grown on surfaces with different stiffness. The cells were stained with Vinculin antibody to visualize FAs and Phalloidin to detect F-actin. Quantification of cell area (median ± 95% confidence interval), F-actin length (mean ± SD) and FA area (median ± 95% confidence interval) were shown in (**E**), (**F**), and (**H**), respectively. Representative images were shown in (**G**). (**I**) Young’s modulus of WT or *PXN*^-/-^ HeLa cells expressing EV, PXN WT, 2SA, or 2SD was measured by AFM. (**J**-**K**) WT or *PXN*^-/-^ HeLa cells reconstituted with EV, PXN WT, 2SA, or 2SD were cultured in 3D collagen gels. The gel size at different time points were photographed (**J**) and quantified (**K**). Data are shown as mean ± SEM from 3 independent experiments. (**L**, **M**) WT or *PXN*^-/-^ HeLa cells reconstituted with EV, PXN WT, 2SA, or 2SD were analyzed by TFM. The total cellular force was quantified in (**M**). ns, not significant; *p < 0.05; **p < 0.01; ***p < 0.001 by one-way ANOVA.

We next asked if PXN phosphorylation by ULK1/2 affect breast cancer cell metastasis *in vivo*. We injected PXN-reconstituted *PXN*^-/-^ MDA-MB-231 cells, which also stably expressed luciferase, into nude mice through their tail vein. Bioluminescence imaging using luciferase showed that compared to WT PXN, cells harboring the PXN 2SA mutant metastasized much more extensively to the lung, whereas the 2SD mutant decreased lung metastases (Fig. 4C–4D). These results suggest that ULK1/2-dependent phosphorylation of PXN restrains breast cancer metastasis.

To test the consequences of PXN phosphorylation in mechanotransduction, we used *PXN*^-/-^ HeLa cells, which exhibited severely compromised ability to spread, rearrange their F-actin cytoskeletal network, and assemble FAs on substrates of high stiffness. WT PXN potently rescued these defects of *PXN*^-/-^ cells. In contrast, expression of 2SA PXN caused the cells to over-spread, arrange a more elaborate actin network, and assemble more FAs, (Fig. 4E–4H and S4E–4F). Next, we sought to investigate if PXN phosphorylation alters actin assembly dynamics. We used cytochalasin D to disrupt intracellular actin polymerization and then removed the inhibitor to permit actin growth over time. This experiment showed that F-actin polymerized more quickly in cells expressing the 2SA mutant than those with either WT or 2SD PXN (Fig. S4G–S4H). This observation suggests that ULK1/2 limits cell spreading and actin remodeling through phosphorylating PXN in response to mechanical inputs. The activity of Rho GTPases family, including RhoA and Rac1, is tightly associated with cell mechanotransduction (Lawson and Burridge, 2014; Ohashi et al., 2017). For instance, RhoA promotes the formation of actin stress fibers and the production of contractile forces and Rac1 facilitates FA maturation (Lawson and Burridge, 2014; Ohashi et al., 2017). Therefore, we reasoned that PXN phosphorylation alters the overall cellular activity of RhoA and Rac1. We performed GST pull-down assay with GST-RTNK and GST-PAK1, which specifically bind to activated RhoA and Rac1, respectively. Interestingly, *PXN*^-/-^ cells showed low levels of active RhoA and Rac1, which were greatly increased by reconstituting WT PXN. Expression of PXN 2SA resulted in much higher degree of RhoA and Rac1 activation (Fig. S4I–4J), entirely consistent with the enhanced mechanotransduction of these cells. Furthermore, we used AFM to measure the impact of PXN phosphorylation on cellular stiffness. Depleting PXN substantially softened the cells, which were markedly reversed by reintroducing WT PXN (Fig. 4I). Cells expressing phospho-defective 2SA PXN were much stiffer compared to those expression WT or 2SD PXN (Fig. 4I) and exerted significantly augmented contractile forces towards the matrix, as evidenced by both 3D collagen contraction assay (Fig. 4J–4K) and TFM (Fig. 4L–4M). Taken together, these results suggest that phosphorylation of PXN by ULK1/2 suppresses cell mechanics, dysfunction of which facilitates breast cancer metastasis.

### Phosphorylation of PXN by ULK1/2 alters its biophysical properties

FAs are the major cellular structure responsible for mechanosensing and mechanotransduction. The intracellular portion of FAs consists of hundreds of proteins and exhibits liquid-like properties (Horton et al., 2015; Kuo et al., 2011). Intriguingly, several recent investigations have reported a role for liquid-liquid phase separation (LLPS) in governing FA dynamics (Case et al., 2022; Li et al., 2020; Wang et al., 2021; Zhu et al., 2020). Consistent with these reports, our unpublished data suggest that PXN undergoes LLPS to promote the macromolecular assembly of the cytosolic FA complex (Wang et al., 2022). Given these findings, we hypothesized that ULK1/2-mediated phosphorylation of PXN might influence PXN LLPS, and therefore FA dynamics, cancer cell mechanics, and metastasis. To test this hypothesis, we first purified recombinant PXN WT, 2SA, and 2SD from *E. coli* (Fig. S5A) and assessed the behavior of these proteins in vitro. PXN WT began forming micro-sized droplets at a concentration of 6.25 μM in physiologically relevant buffer (150 mM NaCl, pH 7.0) without any molecular crowder (Fig. 5A). The threshold concentration required for LLPS was slightly higher for PXN 2SA (~10 μM) and dramatically higher for PXN 2SD (> 25 μM) (Fig. 5A–5B), suggesting an inhibitory effect of PXN phosphorylation on PXN LLPS.

**Figure 5.**
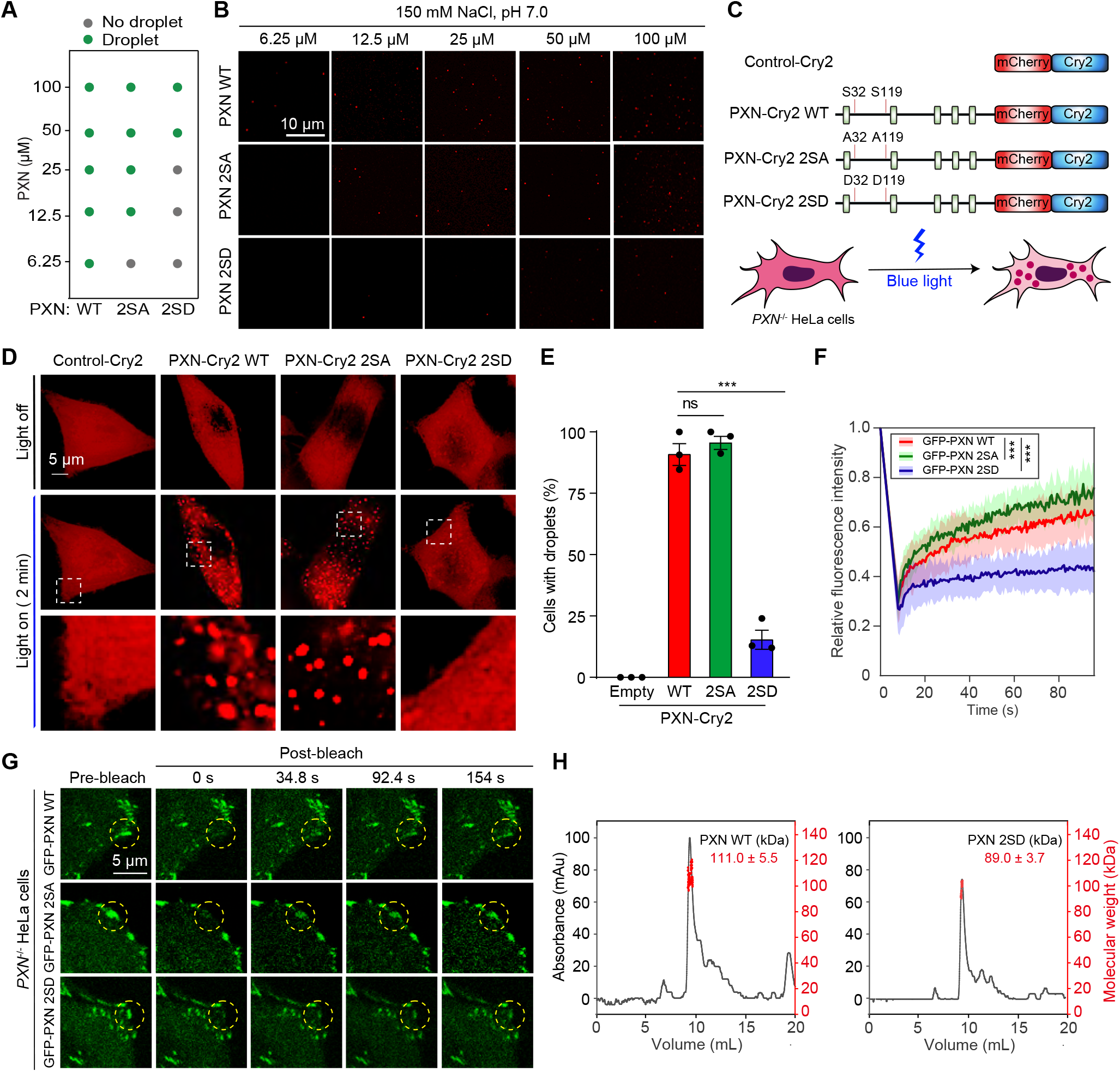
Phosphorylation of PXN by ULK1/2 alters its biophysical properties. (**A**) Phase diagram of PXN WT, 2SA, and 2SD with increasing protein concentrations (**B**) Representative images of LLPS of PXN WT, 2SA, 2SD at different concentrations. (**C**) Schematic illustrations of the Control-Cry2 and PXN-Cry2 WT, 2SA, and 2SD. (**D**, **E**) Representative images of *PXN*^-/-^ HeLa cells stably expressing Control-Cry2 or PXN-Cry2 WT, 2SA, 2SD constructs treated with 2 min of 488 nm blue light. The percentage of cells with Opto-PXN droplet formation was quantified (**E**). Data are shown as mean ± SEM from 3 independent experiments. (**F**, **G**) *PXN*^-/-^ HeLa cells stably expressing GFP-PXN WT, 2SA, or 2SD were subjected to FRAP. The fluorescence intensity was monitored over time and quantified (**G**). Data are shown as mean ± SD. n = 9. (**H**) The molecular weight of recombinant WT or 2SD PXN was determined by multi-angle light scattering. ns, not significant; ***p < 0.001 by one-way ANOVA.

We next sought to determine whether a similar phenomenon occurs in cells by using an optogenetic tool wherein intrinsically disordered regions (IDRs) of target proteins are fused with Cry2 (Shin et al., 2017). In this system, Cry2 oligomerizes upon blue light illumination, serving as the initial nucleation event to trigger intracellular LLPS (Shin et al., 2017). Because the N terminus of PXN was predicted to be largely disordered (data not shown), we expressed this fragment fused with Cry2-mCherry (PXN-Cry2) as well as mCherry-Cry2 alone (Control-Cry2) (Fig. 5C). Whereas *PXN*^-/-^ HeLa cells stably expressing Control-Cry2 remain unresponsive to brief blue light exposure (2 mins), more than 90% of cells expressing PXN-Cry2 WT rapidly formed numerous cytoplasmic condensates upon blue light exposure (Fig. 5D–5E). Similar results were obtained with cells expressing PXN-Cry2 2SA (Fig. 5D–5E). Strikingly, cells expressing PXN-Cry2 2SD were almost refractory to blue light stimulation (Fig. 5D–5E), indicating diminished light-induced PXN LLPS due to PXN phosphorylation.

We next performed fluorescence recovery after photobleaching (FRAP) to assess the molecular dynamism of PXN mutants in cells. All the GFP-PXN fusion protein, including WT, 2SA, and 2SD, localized to the FAs. When we photobleached the GFP-PXN within the FAs, the recovery rate was significantly faster for PXN 2SA and significantly slower for PXN 2SD, compared to GFP-PXN WT, (Fig. 5F–5G), indicating weaker physical interactions between the 2SD mutant with the FA constituents.

Given that high order homotypic oligomerization often promotes protein LLPS (Zhang et al., 2020), we sought to determine if PXN forms oligomers and if so, is it affected by phosphorylation? Using *in vitro* cross-linking with bis-sulfosuccinimidyl suberate (BS^3^), we found that purified PXN WT was capable of forming dimers. This ability was prominently diminished for the PXN 2SD mutant (Fig. S5B). We further took advantage of dynamic light scattering to evaluate the oligomeric status of recombinant WT and 2SD PXN proteins. PXN monomer was predicted to be 60.9 kDa, and the molecular weight of PXN WT in aqueous solution was determined to be 111.0 kDa, suggesting that the majority of PXN WT likely exists as dimers under our experimental condition (Fig. 5H). In contrast, the molecular weight of PXN 2SD was decreased to 89.0 kDa, suggesting a decreased proportion of PXN 2SD dimers in the solution (Fig. 5H). Consistent with this observation, we found that PXN 2SD was less likely to initiate self-interaction in cells as shown by immunoprecipitation (Fig. S5C). Taken together, we propose that phosphorylation of PXN by ULK1/2 weakens homotypic interactions, decelerates molecular thermodynamics, and therefore increases the threshold for LLPS, which subsequently leads to impeded FA assembly/maturation.

### ULK1/2 and FAK act antagonistically to regulate cell mechanotransduction

Notably, the serine phosphorylation sites of PXN mediated by ULK1/2 (S32 and S119) are adjacent to the tyrosine phosphorylation sites mediated by FAK/Src (Y31 and Y118) (Brown and Turner, 2004). All of these residues are evolutionarily conserved in vertebrates (Fig. S6A), suggesting their functional importance. Given the opposing trends of ULK1/2 and FAK/Src expression in breast cancer tissues (Fig. 1A, and Fig. S1A), we hypothesized that these two sets of kinases might have an antagonistic regulatory relationship in mechanotransduction. To test this hypothesis, we generated PXN mutants with altered FAK/Src phosphorylation sites, either alone or in combination with mutated ULK1/2 phosphorylation sites. Thus, we generated forms of PXN that were phospho-defective (PXN 2YF), phospho-mimetic (PXN 2YE), quadruple phospho-defective (PXN 2YF/2SA), or quadruple phospho-mimetic (PXN 2YE/2SD), and then reconstituted *PXN*^-/-^ cells with these proteins (Fig. S6B). As expected, PXN WT rescued the aberrant cell spreading, disrupted actin stress fibers, and defective FA assembly/maturation of the *PXN*^-/-^ cells. Whereas expression of PXN 2YF yielded phenotypes indistinguishable from those associated with PXN WT, expression of PXN 2YE resulted in even greater cell spreading, longer F-actin length, and greater FA assembly than did expression of PXN WT (Fig. 6A–6D), suggesting a positive role for FAK/Src-mediated phosphorylation of PXN in cell mechanotransduction. Importantly, the impact of PXN phosphorylation at Y31 and Y118 on cellular morphology and FA assembly/maturation was reversed by PXN phosphorylation at S32 and S119 (Fig. 6A–6D). Compared to WT PXN, PXN 2YF expression resulted in a small but consist decrease in cell contraction in the 3D collagen contraction assay. This compromised cell contractility due to defective FAK phosphorylation was dramatically rescued by the quadruple phospho-defective (2YF/2SA) PXN mutant (Fig. 6E–6F), indicating functional antagonism of serine and tyrosine phosphorylation of PXN in cell contraction.

**Figure 6.**
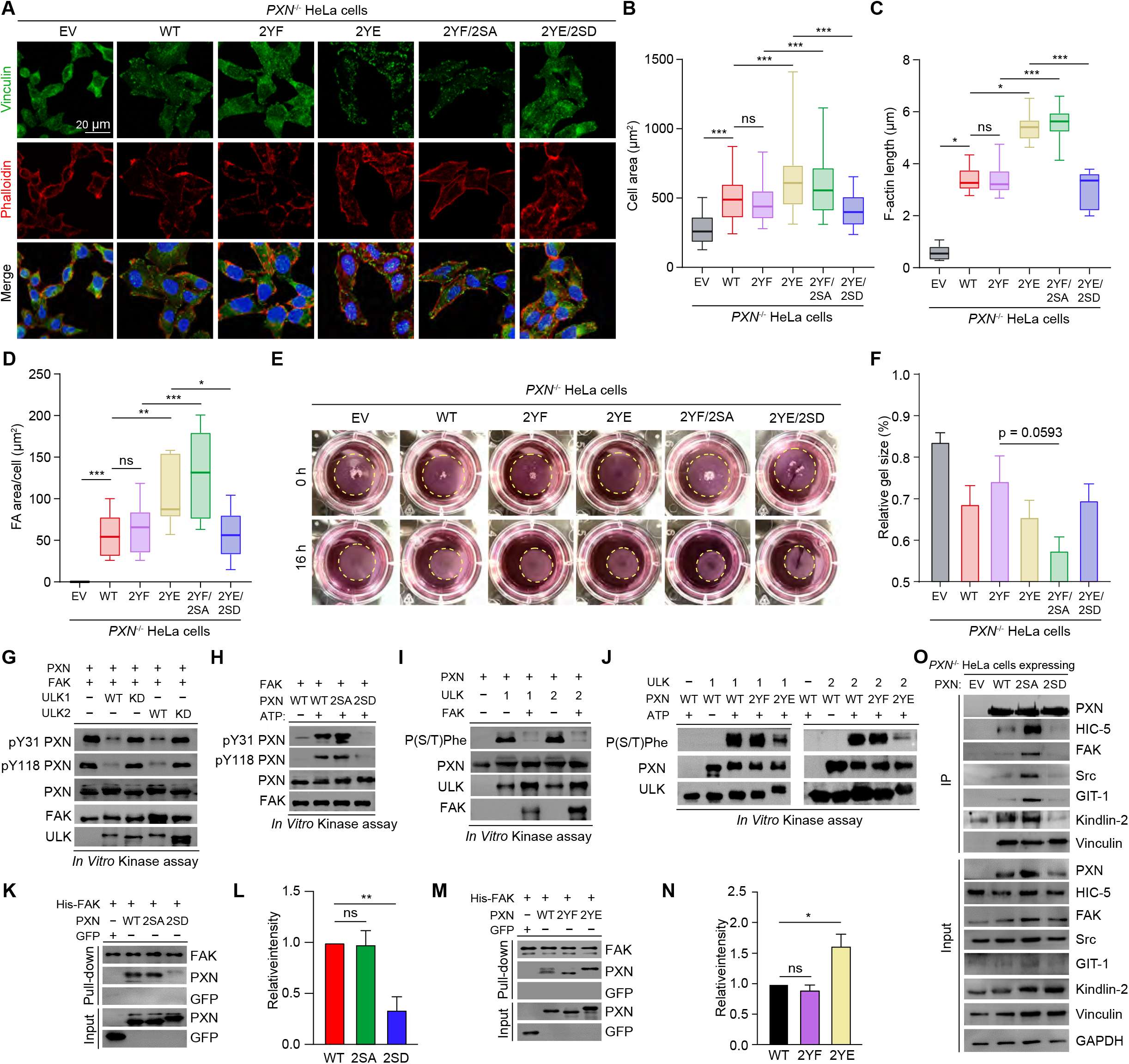
ULK1/2 and FAK act antagonistically to regulate cell mechanotransduction. (**A**-**D**) *PXN*^-/-^ HeLa cells expressing EV, or PXN WT, 2YF, 2YE, 2YF/2SA, 2YE/2SD were stained with Vinculin antibody and Phalloidin. Quantification of cell area, F-actin length and FA area were shown in (**B**), (**C**), and (**D**), respectively. (**E**, **F**) *PXN*^-/-^ HeLa cells expressing EV, or PXN WT, 2YF, 2YE, 2YF/2SA, 2YE/2SD were seeded in 3D collagen gels. The gel size was photographed (**E**) and quantified (**F**). Data are shown as mean ± SEM from 3 independent experiments. (**G**-**J**) *In vitro* kinase assay was performed using the indicated proteins. The samples were analyzed by immunoblot with the indicated antibodies. (**K**-**N**) Pull-down assay using recombinant FAK and PXN mutants. The relative band intensities of PXN were determined by densitometry (**L**) and (**N**). Data are shown as mean ± SEM from 3 independent experiments. (**O**) *PXN*^-/-^ HeLa cells reconstituted with EV or Flag-tagged PXN WT, 2SA, 2SD were subjected to Flag immunoprecipitation. The eluted samples were analyzed by immunoblot with the indicated antibodies. ns, not significant; *p < 0.05; **p < 0.01; ***p < 0.001 by one-way ANOVA.

We next performed biochemical studies to investigate the apparent antagonism between ULK1/2 and FAK/Src. In *in vitro* kinase assays, pre-incubation of PXN with ULK1/2 WT, but not ULK1/2 KD, dramatically inhibited PXN phosphorylation by FAK *in vitro* (Fig. 6G), consistent with our observation that PXN 2SD showed lower levels of phosphorylation by FAK and Src compared with PXN WT (Fig. 6H, S6C). Conversely, pre-incubation of PXN with FAK markedly reduced PXN phosphorylation by ULK1/2 (Fig. 6I), and the PXN 2YE mutant showed decreased levels of phosphorylation by ULK1/2 (Fig. 6J). These data demonstrate that ULK1/2 and FAK/Src competitively phosphorylate PXN.

We further noted that while the PXN 2SD mutant showed decreased interaction with FAK and Src (Fig. 6K–6L), PXN 2YE exhibited enhanced binding to FAK and Src (Fig. 6M–6N, and S6D–S6G). This altered binding affinity of PXN 2YE was specific for FAK, as we detected no changes in binding to other interacting partners, such as Vinculin and Kindlin-2, in the PXN phospho-mutants compared with PXN WT (Fig. S6H–S6K). We next examined the binding partners of PXN 2SA and 2SA by co-immunoprecipitation in cells. We found that PXN 2SA displayed stronger association with several binding partners, including HIC-5, FAK, Src, GIT-1, and kindlin-2, whereas PXN 2SD showed little interaction with these proteins (Fig. 6O). Therefore, we propose that PXN phosphorylation by ULK1/2 disfavors heterotypic interactions between FAK/Src and PXN, and subsequently inhibits PXN tyrosine phosphorylation. This PXN-centric signaling node functions as a dynamic modulator of mechanotransduction, which is linked to the metastatic potential of breast cancer cells.

## Discussion

Here, we demonstrate that independent of autophagy regulation, ULK1/2 negatively govern breast cancer cell mechanotransduction and migration through phosphorylating PXN. Phosphorylation of PXN by ULK1/2 weakens homotypic interactions, decelerates molecular thermodynamics, and therefore increases the threshold for LLPS, which subsequently leads to impeded FA assembly/maturation and aberrant mechanotransduction.

Our demonstration of ULK1/2 in autophagy-independent mechanotransduction adds to their expanding non-canonical functional spectrum (Joo et al., 2016; Wang et al., 2018; Wang and Kundu, 2017; Wang et al., 2019). Although ULK1/2-mediated phosphorylation of PXN in mechanotransduction appears to be unrelated to autophagy under nutrient-rich culture conditions, it does not exclude the possibility that ULK1/2 may regulate autophagy flux through PXN phosphorylation. Indeed, PXN has been implicated in autophagy regulation in several independent studies (Chen et al., 2008; Lv et al., 2022; Sharifi et al., 2016). ULK1/2 may redirect cellular resources to autophagy-related pathways in response to metabolic stress (Wang and Kundu, 2017) and we therefore speculate that such a rewiring mechanism may also apply for the ULK1/2-PXN complex. For example, ULK1/2 might relay mechanical signals to downstream machinery by phosphorylating PXN in response to mechanical stimuli where both AMPK and mTOR are activated, but redirect PXN (and perhaps other FA components) for stress-induced autophagy under starvation conditions, when AMPK is activated and mTOR is inhibited. Thus, our study establishes a ULK1/2-PXN signaling nexus in mechanotransduction in breast tumors, which may function as a molecular switch between mechanotransduction and autophagy regulation.

The macromolecular assembly of FAs in the cytoplasm is facilitated by LLPS. Several lines of evidence suggest that the phase separation properties of FA-associated proteins, and therefore FA dynamics, are modulated by ECM mechanical stimuli (Case et al., 2022; Li et al., 2020; Wang et al., 2021; Zhu et al., 2020). For example, LIMD1, which is structurally similar to PXN, was recently shown to undergo LLPS and mechanical force-dependent localization to mature FAs (Wang et al., 2021). Yet, how the dynamics of FAs are regulated in response to mechanostimulation, and the role of LLPS in this process, are much less clear. To our knowledge, the present study represents the first demonstration of a causal relationship between posttranslational modification and LLPS, FA dynamics, mechanotransduction, and tumor metastasis. Significant efforts are being devoted to targeting the biophysical properties of biomolecular condensates for treating disease. Hence, modulating the material properties of FAs, which are membrane-associated condensates, could provide a new perspective for breast cancer therapy.

Our studies also demonstrate that ULK1/2 and FAK/Src act oppositely in mechanotransduction by competitive phosphorylation. Phosphorylation of PXN by one kinase (ULK1/2 or FAK/Src) substantially attenuated the phosphorylation of PXN by the other (FAK/Src or ULK1/2) both *in vitro* and in cells, and we found that phosphorylation by these kinases acted antagonistically in mechanotransduction. Such a precise regulatory mechanism likely entails an upstream kinase to coordinate the activity of ULK1/2 and FAK/Src such that cells properly respond to mechanical stimulation. The identity of such kinase(s) remains unknown. Importantly, FAK/Src-mediated phosphorylation at Y31 and Y113, and ULK1/2-mediated phosphorylation at the neighboring S32 and S119, are evolutionarily conserved in the vertebrates, highlighting the importance of this signaling nexus. Such an elegant regulatory mechanism by serine and tyrosine kinases reflects the intricate cellular programs to gate-keep mechanosensing and mechanotransduction.

Dysfunctional mechanosensing and mechanotransduction contribute to tumorigenesis and metastasis (Broders-Bondon et al., 2018; Riehl et al., 2020). Non-tumorous cells are sensitive to mechanical cues from the ECM. In contrast, tumorous cells often are refractory to these cues and gain uncontrolled growth and migration. Here, we reveal that breast cancer cells hijack the suppressive forces of mechanotransduction mediated by the ULK1/2-PXN complex and boost the mechanotransduction-promoting mechanisms fulfilled by the FAK/Src-PXN complex to accelerate tumor metastasis. We propose that the ULK1/2-PXN-FAK/Src signaling node in the mechanotransduction pathway can be targeted to treat breast tumors. In fact, FAK inhibitors hold great promise for breast cancer treatment (Lorusso et al., 2022; Timbrell et al., 2021). As ULK1/2 agonists are now commercially available, augmenting ULK1/2 kinase activity may represent a new therapeutic avenue.

## Materials and methods

### Clinical tissue analysis

Clinical study was approved by the Medical Ethics Committee of Zhongshan Hospital Affiliated to Xiamen University in accordance with the Declaration of Helsinki. All breast tissues were obtained from the tissue bank of Zhongshan Hospital (Xiamen University). Case 1 and 3 are luminal A (ER^+^ and PR^+^); Case 2 and 5 are luminal B (HR^+^ and HER-2^+^); Case 4 and 6 are HER-2-enriched (ER^-^, PR^-^, and HER-2^+^). Tissue samples were lysed in ice-cold lysis buffer and subjected to immunoblot analyses.

### *In vivo* tail vein injection

MDA-MB-231 cell lines stably expressing luciferase (1 × 10^6^ cells per mouse) were injected into female nude mice at 6-8 weeks through tail vein. 4 weeks after injection, mice were intraperitoneally injected with 10 mg/ml D-luciferin (Acmec Biochemical; D37330) and imaged using Caliper IVIS Lumina II. Total photon flux was measured. These experiments were performed in accordance with protocol approved by the Animal Care and Use Committee of Xiamen University.

### Plasmids

cDNA was amplified by using a standard PCR-based approach. DNA restriction endonucleases were used to linearize vector backbones and the target fragments were amplified by high-fidelity DNA polymerase 2 × Phanta Max Master Mix (Vazyme; P515-01). 2 × MultiF Seamless Assembly Kit was used to construct the plasmids (Abclonal; RK21020). Sanger sequencing was performed to confirm sequence accuracy.

### Cell culture, transient transfection, lentivirus infection, and drug treatment

The HEK293T (CRL-1573), MDA-MB-231 (HTB-26), HeLa (CCL-2) cell lines were purchased from (American Type Culture Collection; ATCC). Cells were cultured in DMEM (L120KJ) containing 10% fetal bovine serum (Moybio; S450), penicillin/streptomycin (BasalMedia; S110JV), and Glutamax (BasalMedia; S210JV) at 37°C (5% CO_2_).

For transient expression, transfections were performed with Polyethyleneimine Linear (BIOHUB; 78PEI25000) or Lipofectamine 2000 (Thermo Fisher; 11668019) according to the manufacturer’s instructions. Knockdown experiments were performed with Lipofectamine RNAi Max (Life Technologies; 13778075) according to the manufacturer’s instructions. shRNA sequences were as follows: *shULK1* sense 5’-CG CGGTACCTCCAGAGCAA-3’; *shULK2* sense 5’-CCA GTTCCTACTCAAATAAC-3’; *shFIP200* sense; sh*ATG7* sense 5’-GCTATTGGAACACTGTATAAC-3’; *shATG13* sense: 5’-GAGAAGAATGTCCGAGAAT-3’; *shATG14* sense 5’-GGGAGAGGTTTATCGACAAGA-3’. The siRNAs synthesized by GenePharma (Shanghai, China) were used as follows: si*ULK1*: 5’-CGCGGUACCUCCAGAGCAATT-3’ and 5’-UUGCUCUGGAGGUACCGCGTT-3’; si*FIP200*: 5’-CUGGGACGGAUACAAAUCCAA-3’ and 5’-UUGGAUUUGUAUCCGUCCCAG-3’; si*ATG7*: 5’-GGAGUCACAGCUCUUCCUUTT-3’ and 5’-AAGGAAGAGCUGUGACUCCTT-3’; *siATG13:* 5’-CCAUGUGUGUGGAGAUUUCACUUAA-3’ and 5’-UUAAGUGAAAUCUCCACACACAUGG-3’; *siATG14:* 5’-GGCAAAUCUUCGACGAUCCCAUAUA-3’ and 5’-UAUAUGGGAUCGUCGAAGAUUUGCC-3’,

All shRNAs were constructed using pLKO.1 retroviral vector; inducible ULK1 WT/KD and ULK2 WT/KD were constructed in pCW57.1 retroviral vector. To generate MDA-MB-231 and HeLa cells stably expressing WT or mutant forms of PXN, lentivirus was produced by co-transfecting 293T cells with psPAX2 (Addgene; #12260), pMD2.G (Addgene; #12259) and pLV retroviral vectors containing different *PXN* cDNAs. Supernatants were harvested at 48 h and 60 h and centrifuged at 8000 rpm for 3 min (RT), and then filtered with a 0.22 μm filter. Polybrene (Santa Cruz Technology; sc-134220) was added to facilitate infection. The transduced cells were FACS-sorted by the presence of GFP or selected with antibiotics.

All the chemicals were dissolved in DMSO. SBI-0206965 (20 mM, 18 h), blebbistatin (20 mM, 2 h), Rho activator II (1 mg/mL, 4 h), and Cytochalasin D (5 μM, 3 h) were directly added to the medium.

### Generation of knock-out cell lines with CRISPR/Cas9

PXN knockout cells were created through the CRISPR/Cas9 technology. The guide RNA sequences were designed using online tool the Optimized CRISPR Design (https://portals.broadinstitute.org/gppx/crispick/public). The guide sequence was 5’-ATCCCGGAACTTCTTCGAGC-3’ for human PXN, for 5’-GCCAAGTCTCAGACGCTGCT-3’ human ULK1, for 5’-ATCTTCCAACCTGTTAGCCT-3’ human ULK2. Cells were transiently transfected with PX459 (Addgene; #48139) and selected with puromycin for 2 days. The pool was scattered into a 10 cm petri dish. Single clones were picked up and expanded for sequencing and immunoblotting analyses.

### Immunoblotting

Cells were harvested and lysed with ice cold lysis buffer (20 mM Tris-HCl, pH 7.5, 150 mM NaCl, 1 mM EDTA, 1 mM EGTA, 1% Triton X-100, 2.5 mM sodium pyrophosphate, 1 mM β-glycerolphosphate, protease inhibitor cocktail). The lysates were then centrifuged to clear cell debris. The supernatant was then prepared with an equal amount of 2 × SDS sample buffer, and electrophoretically separated on SDS-PAGE gels. Proteins were transferred to PVDF membranes. After blocking with 5% skin milk, the membranes were probed with the following primary antibodies: rabbit anti-phospho-PXN (Y31) (Invitrogen; 2024882), rabbit anti-phospho-PXN (Y118) (Cell Signaling Technology; 2541S), rabbit anti-PXN (Proteintech; 10029-1-Ig), rabbit anti-PXN (Abcam; ab32084), rabbit anti-α-tubulin (Cell Signaling Technology; 2125S), rabbit anti-FAK (Cell Signaling Technology; 3285S), rabbit anti-phospho-FAK (Y397) (Abcam; ab81298), rabbit anti-phospho-FAK (Y925) (Abcam; ab38512), mouse anti-Flag (GNI; GNI4110-FG), rabbit anti-Vinculin (Proteintech; 26520-1-AP), mouse anti-GAPDH (Santa; sc-32233), rabbit anti-Kindlin-2 (Proteintech; 11453-1-AP), rabbit anti-HIC-5 (Proteintech; 10565-1-AP), mouse anti-HIC-5 (BD Transduction; 611164), mouse anti-β-actin (Proteintech; 66009-1-Ig), mouse anti-GIT1 (BD Transduction; 611396), rabbit anti-Src (Cell Signaling; 2109S), rabbit anti-phospho-Src (Y416) (Cell Signaling; 59548S), rabbit anti-phospho-Src (Y527) ( Cell Signaling; 2105T), mouse anti-Integrin β1 (Abcam; ab30394), mouse anti-Integrin α2β1 (Abcam; ab24697), rabbit anti-α-smooth muscle Actin (Abcam; ab5694), rabbit anti-phospho-Myosin II (S19) (Cell Signaling; 3675S), rabbit anti-ATG5 (Abclonal; A19677), rabbit anti-ATG7(Abclonal; A19604), rabbit anti-ATG13 (Proteintech; 18258-1-AP), rabbit anti-ATG14 (Proteintech; 24412-1-AP), rabbit anti-phospho-ATG14 (S29) (Cell Signaling; 92340S), mouse anti-Myc-Tag (Abclonal; AE010), rabbit anti-FIP200 (Abclonal; A14685), rabbit anti-P62 (Cell Signaling; 48768T), rabbit anti-ULK1 (Cell Signaling; 8054S), Rabbit anti-LC3B(Cell Signaling; 3868S), rabbit anti-Phospho-Phe (Ser/Thr) ( Cell Signaling; 9631S), mouse anti-mTOR (Proteintech; 66888-1-Ig), rabbit anti-AMPKα (Proteintech; 10929-2-AP). The membranes were then incubated with HRP-conjugated secondary antibodies (Jackson ImmunoResearch; 115-035-003, 111-035-003), and bands were detected using chemiluminescence detection kit (Merck Millipore; WBKLS0050).

### Immunoprecipitation and pull-down

Cells were lysed by lysis buffer (20 mM Tris-HCl, pH 7.5, 150 mM NaCl, 1% Triton X-100, 1 mM EDTA, 1 mM EGTA, 1 mM β-glycerolphosphate, 2.5 mM Sodium pyrophosphate, 1 μM leupeptin, 2 mM Na3VO4 and 1 mM phenylmethylsulfonyl fluoride) after washed by PBS. Lysates were centrifuged at 10,000 rpm for 10 min at 4°C, and supernatants were incubated with anti-DYKDDDDK G1 Affinity beads (Genscript; L00432-10) or anti-GFP magnetic beads (Bio-Linkedin; L-1016A) for 3 h at 4°C. Beads were washed 3 times with lysis buffer and the immunoprecipitates were eluted with 2 x SDS sample buffer. SDS elutions were analyzed by western blotting.

His_6_-tagged proteins (His-FAK/Src/Vinculin/Kindlin-2) were first incubated with 20 μL Ni-NTA agarose (Nuptec; NRPB57L-100) in pull-down buffer (50 mM HEPES pH 7.5, 150 mM NaCl, 1mM DTT and 0.1% Triton-100x) at 4°C for 3 h. Then the beads were centrifuged and the supernatants were discarded. After that, 4 μg proteins without the His tag were added and incubated for another 30 min. After washing 2 times with pull-down buffer, bound proteins were denatured with 2 x SDS sample buffer, and subjected to western blotting.

### *In vitro* kinase assay

Flag-tagged ULK1/2 kinases were derived from transfected 293T cells by anti-DYKDDDDK G1 Affinity beads. 5 μg purified PXN and 2 μg kinase were added to the *in vitro* kinase assay buffer (50 mM Tris-HCl, pH 7.6, 1 mM dithiothreitol, 100 mM MgCl_2_, 1 mM ATP) and incubate for 30 min at 37°C. The samples were subjected to western blotting.

### Immunofluorescence

Cells seeded on glass coverslips were fixed with 4% PFA (Leagene; DF0135) at RT for 10 min, permeabilized with 0.1% Triton X-100 (diluted in PBS), then blocked with 3% BSA, and incubated with primary antibodies overnight at 4°C. The following primary antibodies were used: mouse anti-Flag (GNI; GNI4110-FG), rabbit anti-Vinculin (Proteintech; 26520-1-AP), mouse anti-HA (Santa Cruz; sc-7392), and rabbit anti-PXN (Abcam; ab32084). The cells were then incubated with Alexa-Fluor-conjugated secondary antibody (Jackson ImmunoResearch; 115-545-003, 115-585-003) for 1 hour at RT in the dark, and mounted on slides with mounting media (SouthernBiotech; 0100-01). Samples were imaged by Zeiss LSM 900 confocal microscopy with a 63x oil objective.

### Protein expression and purification

For protein expression, plasmids were transformed into BL21(DE3) *E. coli* cells (AngYu; G6030-10). A single colony was inoculated into LB media containing ampicillin and grown in LB media to an optical density of 0.6-0.8 at 37°C, followed by overnight induction with 1 mM isopropyl-b-D-thio-galactopyranoside (IPTG) at 16°C. Cells were pelleted and resuspended in binding buffer (20 mM Tris pH 7.5, 500 mM NaCl) and lysed using a homogenizer. The lysates were cleared by centrifugation and purified by Ni-NTA agarose (Nuptec; NRPB57L-100). The proteins were eluted with elution buffer (20 mM Tris pH 7.5, 500 mM NaCl, 300 mM imidazole). Eluate was concentrated and cleaved with TEV protease at 4°C. The cleaved tags and TEV protease were separated from the protein samples by a second round of Ni-NTA affinity chromatography followed by a Sephadex 200 size-exclusion column in 20 mM Hepes, 100 mM NaCl, pH 8.0. The collected fractions were then verified with SDS–PAGE. Purified fractions were pooled, concentrated, and flash-frozen in liquid nitrogen. FAK purification has been described previously (Wang et al., 2022).

### Gel contraction assay

6 × 10^5^ cells were resuspended in 750 μL medium and added to 200 μL rat tail tendon collagen type I (Shengyou; 200110-50), followed by intensive mixing. The cells and collagen mixture were then seeded into a 6-well plate and cultured at 37°C with 5% CO2 for 30 min. The collagen gels were isolated from the well by shaking gently and then 2 mL medium was added into each well. Photographs of the collagen gels were taken at 0, 16, 32 and 48 h. Fiji was used to measure the area of collagen gels at each time point.

### Atomic force microscopy

AFM experiments were carried out in a Bruker Nanowizard 4 (JPK) mounted onto an inverted optical microscope (Zeiss Observer 7; Zeiss, Germany). Force indentation measurements were carried out using in-house-prepared AFM colloidal probes (a spherical silica bead with a diameter of 10 μm glued on the cantilever by epoxy). Before each experiment, we performed a calibration to determine the elastic coefficient of the cantilever. During the experiment, cells were kept at 37°C in 1 x PBS buffer. Indentations were performed at a loading force of 0.5 nN and a constant speed of 4 μm/s. Young’s modulus was obtained by fitting the force-distance curve to the Hertzian sphere model.

### Preparation of PA gels

Preparation of PA gels with different stiffness was done as previously described (Denisin and Pruitt, 2016). The acrylamide mix was dropped on the glass slides. PA gels were removed from the slide glass after solidification. The stiffness of PA gels was measured by AFM.

We functionalized the PA gels with fibronectin according to the following procedure: (1) PA gels were treated with soaking buffer (137 mM NaCl, 5% glycerol) for 1 hour; (2) After removing the soaking buffer, the PA gels were further treated with buffer mix containing EDC/NHS and conjugation buffer (0.2 M MES, 10% glycerol, pH 4.5) in the dark; (3) The gels were coated with fibronectin at a final concentration of 50 μg/mL at 4°C overnight.

### Traction force microscopy

We manufactured a glass slide with a chamber, and spread PA gel (8 kPa) embedded with 580/605 100 nm-diameter on the bottom of the chamber. The cells were allowed to adhere to the substrate for at least 12 h prior to imaging. Fluorescence images of the embedded beads were captured on LSM 980 (*Zeiss*) confocal imaging system. After collecting the images of the beads, 0.25% trypsin/EDTA was added to detach the cells for at least 5 min, and images of the bead position without cellular forces were captured. Displacement of the beads and reconstruction of force field were calculated based on Matlab algorithm (https://github.com/DanuserLab/TFM) The corresponding gel deformations were obtained by 2D Gaussian distribution interpolation. Using an algorithm based on Fourier transform, the stress field of the gel deformation was calculated.

### Transwell assay

3 × 10^4^ cells were seeded into the upper chamber of the transwell plates (Corning Incorporated; 3422) in FBS-free medium. Medium containing 10% FBS were added to the lower compartment as chemoattractant. After incubation at 37°C for 16-18 h, the migrated cells were fixed by 4% PFA (Leagene; DF0135) for 10 min and stained with 0.2% (w/v) crystal violet (Sangon Biotech; A600331-0100) for 10 min and washed with ddH2O. The stained cells were imaged by Olympus IX51 4 x objective, and Fiji was used to count the cell number. All migration assays were repeated at least 3 times.

### Fluorescence recovery after photo-bleaching

FRAP experiments were performed using water lens on a Cell Discoverer 7 (*Zeiss*) system. Cells were maintained at 37°C and 5% CO2. Cells were imaged for a total of 100 s at a maximum rate of 1 frame per s. After 3 pre-bleach frames, the fluorescence signal from a region of interest (2 μm × 2 μm) was bleached with a 488 nm laser at 100% power. The signal from the bleached ROI was measured for 90 s afterwards. We used ZEN to derive the raw fluorescence values. EasyFRAP software (https://easyfrap.vmnet.upatras.gr/) was used for rectify and normalize the data.

### Protein labelling

The Fluro 488 NHS ester (AAT Bioquest; 1810) or Cy3 NHS ester (AAT Bioquest; 271) were dissolved in DMSO at a concentration of 10 mg/mL. The labelling dye was incubated with the target protein 1:1 (molar ratio) at RT for 1 hour with continuous stirring. Free dye was removed using a desalting column (Merck Millipore; UFC501024), the labeled proteins was aliquoted and flash-frozen in liquid nitrogen.

### *In vitro* LLPS assay

Mixtures in a total solution volume of 2 μL were placed in on slides with double-sided tape. Cover glasses were placed on top to seal the slides. Droplet formation of purified protein was monitored by fluorescence microscopy using a confocal microscope (Zeiss LSM 900).

### Size-exclusion chromatography coupled with multi-angle light scattering (SEC-MALS)

SEC-MALS experiments were carried out on DAWN HELEOS-II (Wyatt Technology Corporation) coupled with a Shimadzu liquid chromatography system equipped with a Superdex 200 10/300 GL (GE Healthcare) gel-filtration column. Protein samples for this experiment were dialyzed to a buffer containing 20 mM Tris (pH 7.2), 400 mM NaCl, 2 mM DTT, and 0.03% NaN3. 200 μL proteins at a concentration of 0.02-0.05 mM were injected and run at a flow rate of 0.03 mL/min. The ASTRA version 6.0.5 (Wyatt Technology Corporation) was used for data collection and analyses.

### Quantification and statistical analyses

Measurements of adhesion area were done with the Imaris software (Bitplane) using the surface reconstruction tool.

To quantify protein colocalization, the cloco2 plug-in (Analyze-Colocalization-Coloc 2) of the image j software was used to calculate the Pearson correlation coefficient between the two target channels per the instructions.

Actin filament were analyzed by applying a steerable filter approach with a deposited software FSegment (http://www2.medizin.uni-greifswald.de/anatomie/forschung/niere/fsegment/). We counted the total length of microfilaments in each image, and then calculated the average length of each cell.

All analyses were performed blindly. All quantitative data are shown as mean ± SEM or median ± 95% confidence interval from n ≥ 3 biological replicates unless otherwise specified. Statistical significance was determined by Student’s t-test or ANOVA as appropriate, and *p < 0.05 was considered statistically significant. Statistical parameters are also reported in the figures and legends.

## Acknowledgements

We are grateful to Dr. Shuguo Sun (Huazhong University of Science and Technology), and Dawang Zhou (Xiamen University) for providing plasmids. We thank Dr. Natalia Nedelsky (St. Jude Children’s Research Hospital) for editing the manuscript. All imaging data were acquired in the Core Facility of Biomedical Sciences, Xiamen University. This work was supported by the National Natural Science Foundation of China (32071235 to B.W.) and Guangdong Basic and Applied Basic Research Foundation (2020A1515111186 to B.W.).

## Figure Legends

**Figure S1.**
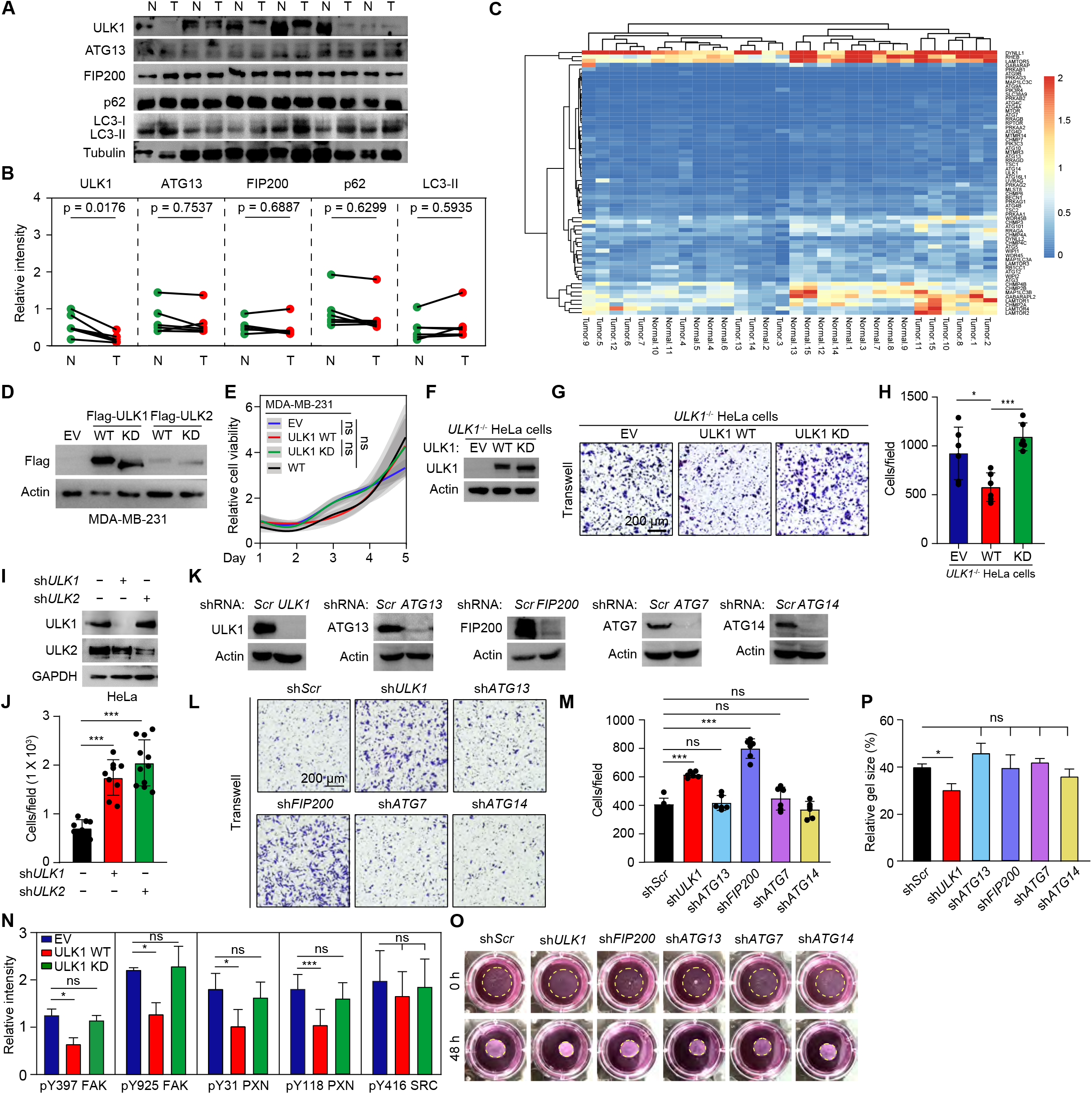
Related to Figure 1. (**A**, **B**) Lysates prepared from normal (N) and cancerous (T) breast tissues were analyzed by immunoblot with the indicated antibodies. The relative band intensities were quantified by densitometry (**B**). Data are presented as mean ± SEM from 6 individual patients. Statistic difference was calculated by paired Student’s t-test. (**C**) mRNA expression levels of key genes in the autophagy pathway determined by sc-RNAseq from normal and malignant breast tissues. (**D**) Cell lysates prepared from WT MDA-MB-231 cells overexpressing EV, flag-tagged ULK1 WT, KD, ULK2 WT or KD were analyzed with immunoblot. (**E**) Growth curves of MDA-MB-231 cells overexpressing EV, flag-tagged ULK1 WT or KD were determined by CCK8 assays. The experiments were repeated 2 times with similar trends. (**F**) Lysates prepared from *ULKT*^-/-^ HeLa cells expressing EV, ULK1 WT or KD were analyzed by immunoblot. (**G**, **H**) *ULK1*^-/-^ HeLa cells expressing EV, ULK1 WT or KD were subjected to transwell assay. The cells migrated to the lower chamber were stained with crystal violet (**G**). Quantification of the cell number was show in (**H**). Data are presented as mean ± SEM. (**I**) Lysate prepared from WT HeLa cells infected with shRNA against scramble (*Scr*), *ULK1, ATG13, FIP200, ATG7*, or *ATG14* were analyzed by immunoblot using the indicated antibodies. (**J**, **K**) WT HeLa cells infected with shRNA against scramble (*Scr*), *ULK1, ATG13, FIP200, ATG7*, or *ATG14* were subjected to transwell assays. Representative photographs of cells stained with crystal violet were shown in (**J**). The number of cells was quantified in (**K**). Data are presented as mean ± SEM. (**L**) HeLa cells infected with shRNA against scramble (*Scr*), *ULK1*, or *ULK2* were analyzed by immunoblot. (**M**) Quantification of cell number of sh*Scr*, *shULK1*, or *shULK2* infected HeLa cells in the transwell assay. Data are presented as mean ± SEM. (**N**) The relative band intensities from Figure 1H were quantified by densitometry. Data are presented as mean ± SEM from 3 independent experiments. (**O**, **P**) HeLa cells infected with shRNA against scramble (*Scr*), *ULK1, ATG13, FIP200, ATG7*, or *ATG14* were cultured in 3D collagen gels. The gel size was photographed (**O**) and quantified (**P**). Data are shown as mean ± SEM from 3 independent experiments. ns, not significant; *p < 0.05; ***p < 0.001 by one-way ANOVA.

**Figure S2.**
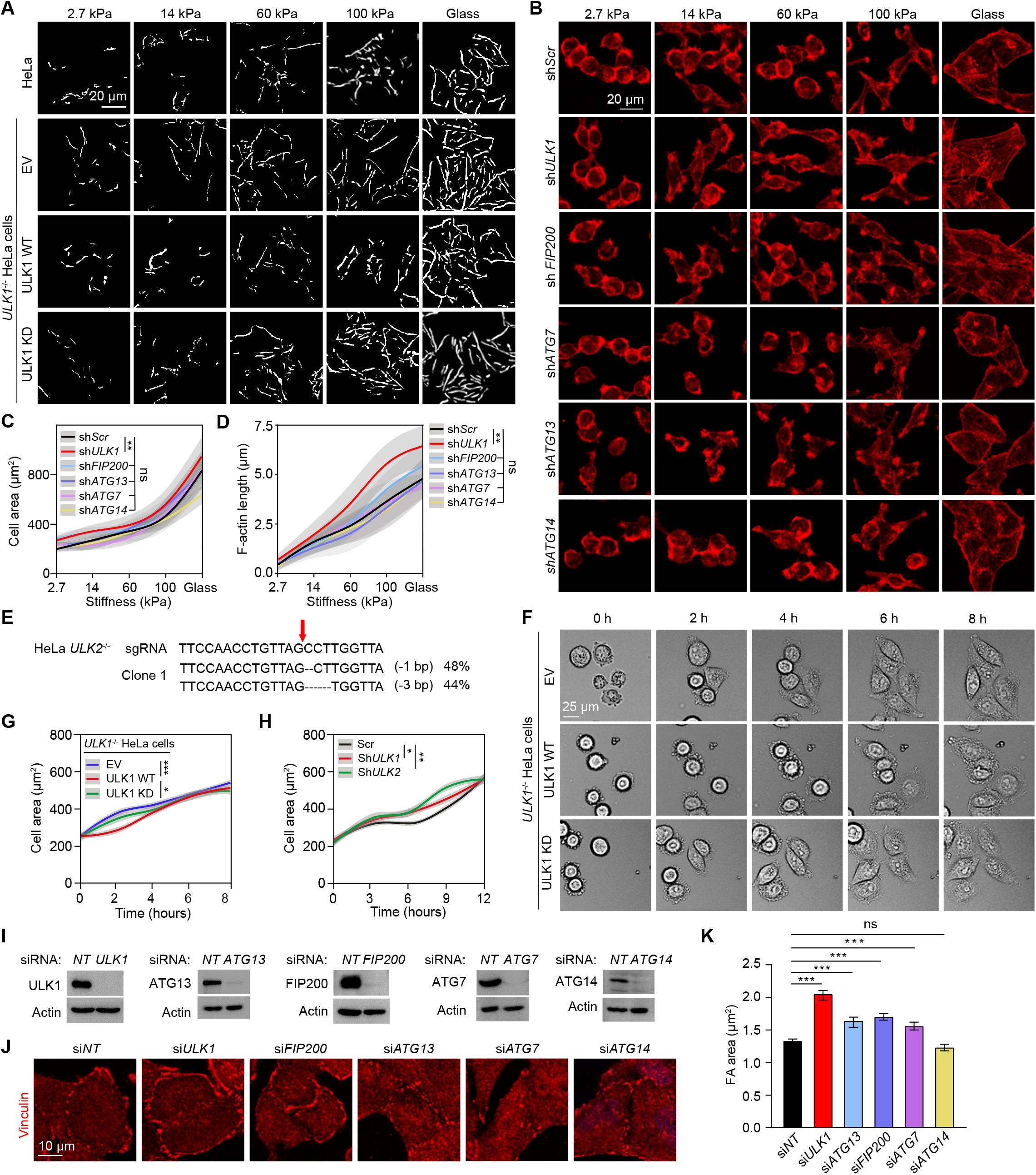
Related to Figure 2. (**A**) Representative masks of Phalloidin staining from Figure 2C. (**B**-**D**) HeLa cells transfected with shRNA against scramble, *ULK1, FIP200, ATG7, ATG13, ATG14* were cultured on surfaces with different stiffness. The cells were stained with Phalloidin (**B**). The cell area (median ± 95% confidence interval) and F-actin length (mean ± SD) were quantified and shown in (**C**) and (**D**), respectively. (**E**) Genomic sequence of *ULK2*^-/-^ HeLa cells. The arrow indicates the cleavage site by Cas9. The numbers listed on the right denote allele frequency. (**F**, **G**) *ULK1*^-/-^ HeLa cells expressing EV, ULK1 WT or KD were subjected to live cell imaging to monitor cell spreading. Representative DIC images at different time points after cell seeding were shown in (**F**). The cell area (median ± 95% confidence interval) was quantified (**G**). (**H**) HeLa cells transfected with shRNA against scramble, *ULK1*, or *ULK2* were subjected to live cell imaging to monitor cell spreading. The cell area (median ± 95% confidence interval) was quantified. (**I**). Cell lysates prepared from HeLa cells transfected with siRNA against non-target (NT), *ULK1, FIP200, ATG7, ATG13, ATG14* were analyzed with immunoblot using the indicated antibodies. (**J**, **K**) HeLa cells transfected with siRNA against NT, *ULK1, FIP200, ATG7, ATG13*, or *ATG14* were fixed and immunostained with antibodies against Vinculin (**J**). FA size was quantified in (**K**). Data are presented as median ± 95% confidence interval. ns, not significant; *p < 0.05; **p < 0.01; ***p < 0.001 by one-way ANOVA.

**Figure S3.**
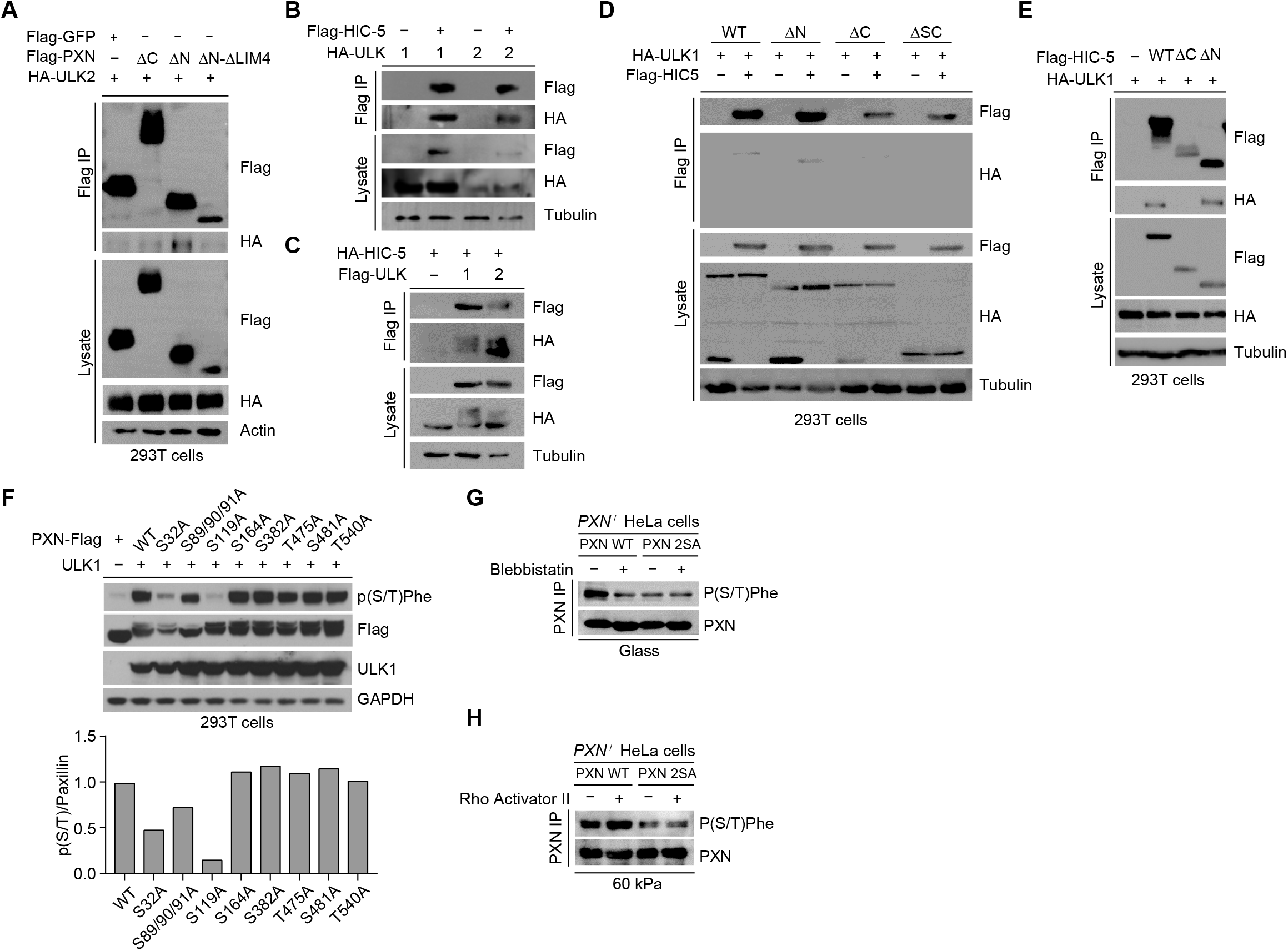
Related to Figure 3. (**A**-**E**) 293T cells transfected with the indicated constructs were subjected to Flag immunoprecipitation. The eluted samples were analyzed by immunoblot using the indicated antibodies. (**F**) Lysates prepared from 293T cells transfected with the indicated constructs were analyzed by immunoblot with the indicated antibodies (top). The relative band intensities of P(S/T) were quantified by densitometry and plotted (bottom). (**G**) *PXN*^-/-^ HeLa cells reconstituted with PXN WT or 2SA growing on glass were treated with DMSO or blebbistatin. PXN was immunoprecipitated and analyzed by immunoblot with P(S/T) antibody. (**H**) *PXN*^-/-^ HeLa cells reconstituted with PXN WT or 2SA growing on 60 kPa gel were treated with DMSO or Rho activator II. PXN was immunoprecipitated and analyzed by immunoblot with P(S/T) antibody.

**Figure S4.**
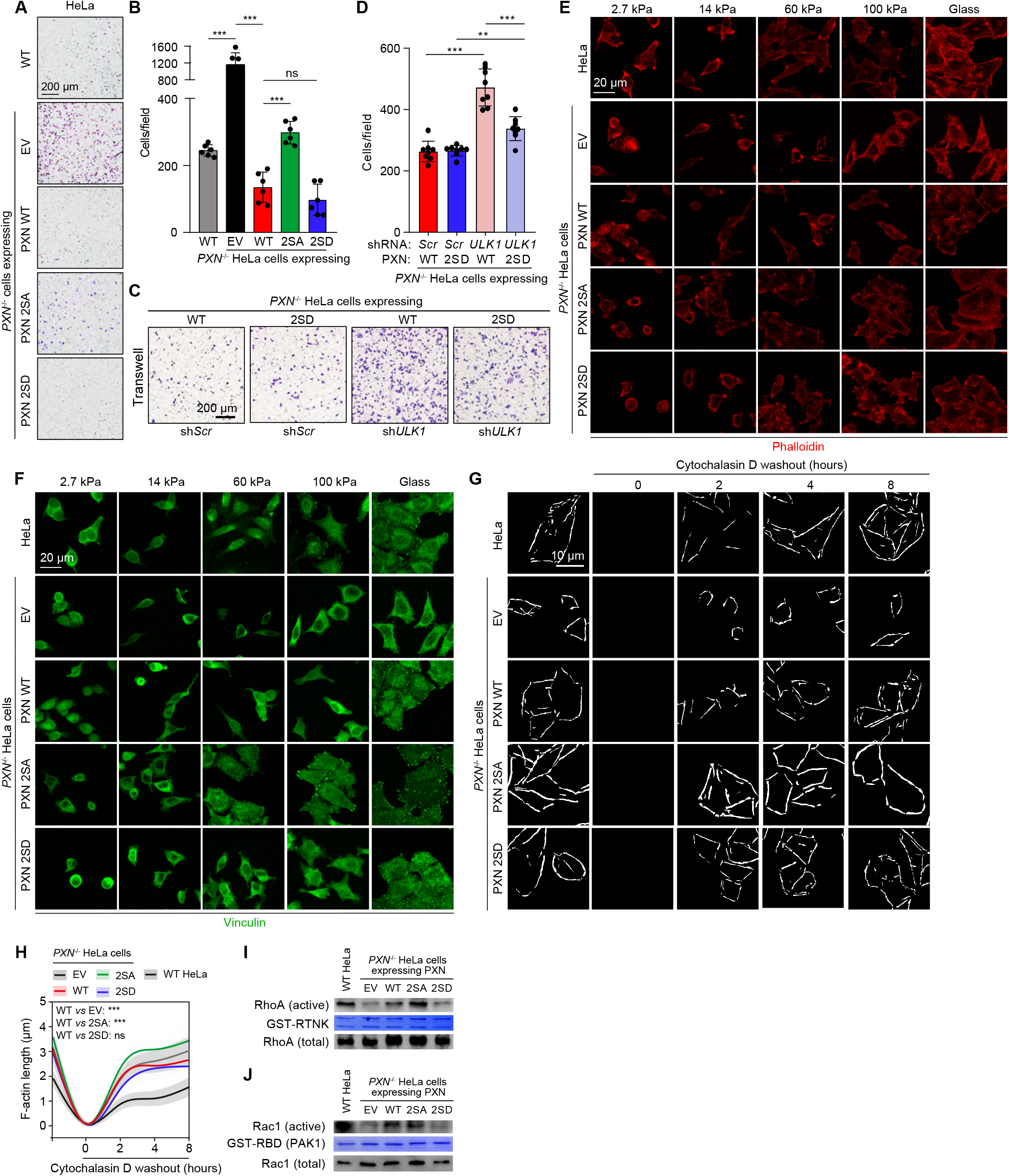
Related to Figure 4. (**A**, **B**) WT or *PXN*^-/-^ HeLa cells reconstituted with EV, PXN WT, 2SA, or 2SD were subjected to transwell assay. The cells were stained with crystal violet (**A**). Quantification of cells migrated to the lower chambers was show in (**B**). Data are presented as mean ± SEM. (**C**, **D**) *PXN*^-/-^ HeLa cells reconstituted with PXN WT or 2SD were transfected with shRNA against scramble or *ULK1*. These cells were then subjected to transwell assay. The cells migrated to the lower chambers were stained with crystal violet (**C**) and quantified (**D**). Data are presented as mean ± SEM. (**E**, **F**) WT or *PXN*^-/-^ HeLa cells expressing EV, PXN WT, 2SA, or 2SD grown on surfaces with different stiffness were stained with Phalloidin to detect F-actin (**E**) and Vinculin antibody to visualize FAs (**F**). (**G**, **H**) WT or *PXN*^-/-^ HeLa cells expressing EV, PXN WT, 2SA, or 2SD grown on glass were treated with Cytochalasin D. The cells were fixed at different time points after Cytochalasin D washout, and visualized with Phalloidin, the masks of which were shown in (**G**). F-actin length (mean ± SD) were quantified. (**I**, **J**) Lysates were collected from *PXN”* HeLa cells expressing EV, PXN WT, 2SA, or 2SD and incubated with RTNK (**I**) or GST-PAK_70–106_ (GST-PAK) (**J**). The levels of total and active RhoA (RTNK bound) or active Rac1(GST-PAK bound) were then analyzed by coomassie blue staining and immunoblotting, respectively. ns, not significant; **p < 0.01; ***p < 0.001 by one-way ANOVA.

**Figure S5.**
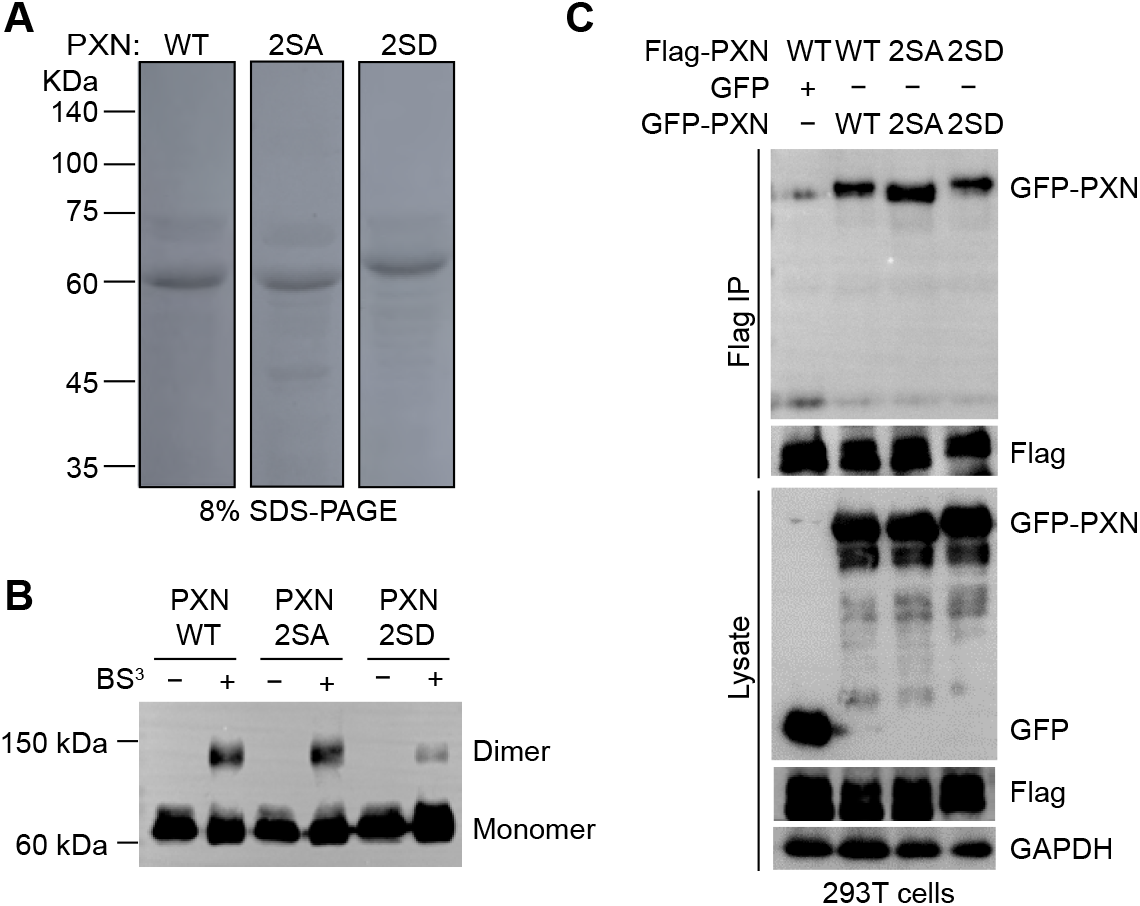
Related to Figure 5. (**A**) Recombinant PXN proteins were separated by 8% SDS-PAGE gels, and visualized by Coomassie Blue. (**B**) PXN WT, 2SA, 2SD were overexpressed in the 293T cells, immunoprecipitated, and subjected to BS^3^ treatment. The crosslinked PXN samples were separated by SDS-PAGE electrophoresis and detected by PXN antibody. (**C**) 293T cells transfected with the indicated constructs were subjected to Flag immunoprecipitation. The eluted samples were analyzed by immunoblot using the indicated antibodies.

**Figure S6.**
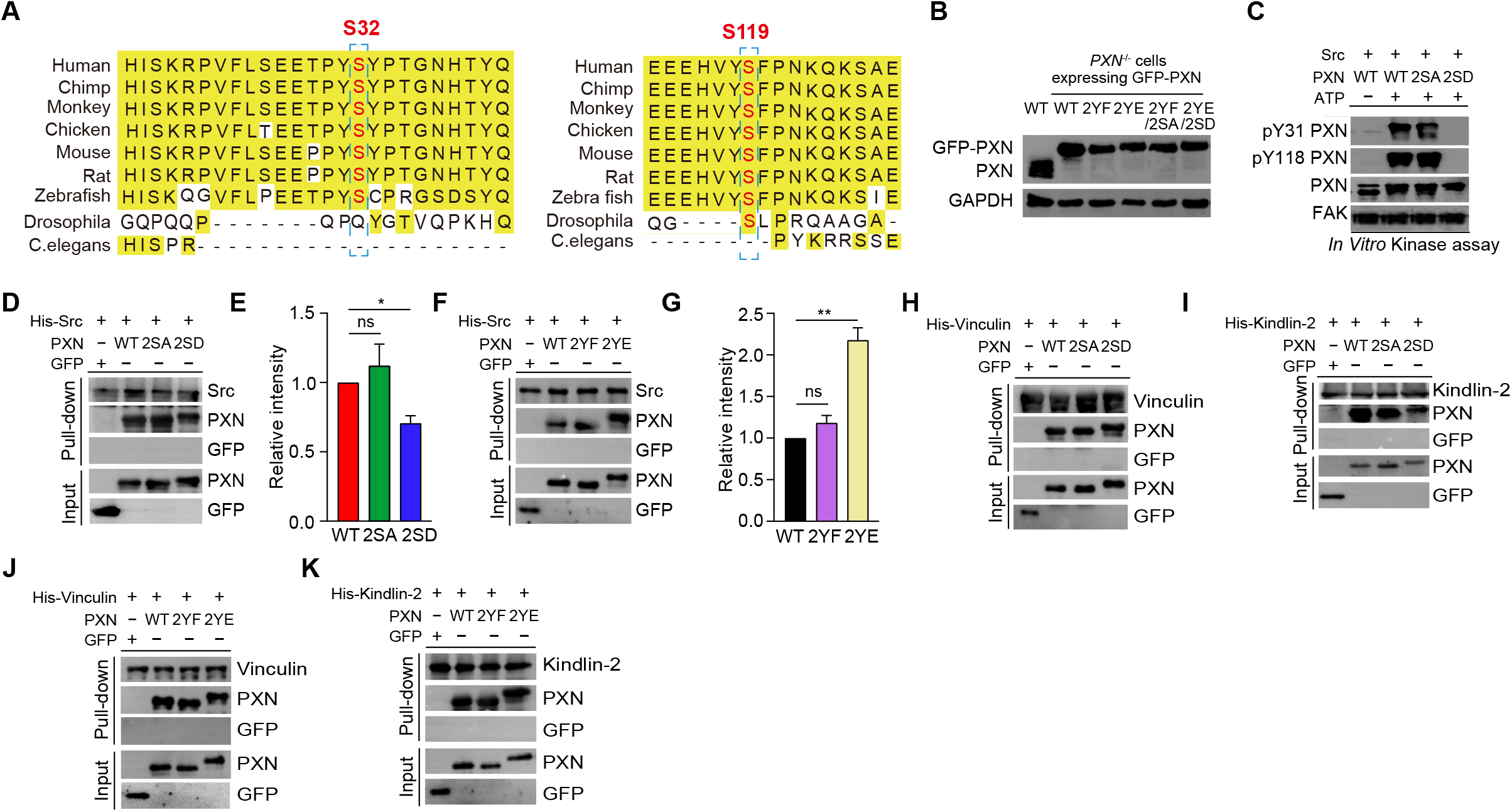
Related to Figure 6. (**A**) Sequence alignment of PXN orthologs of different species. (**B**) *PXN*^-/-^ HeLa cells stably expressing GFP-tagged PXN WT, 2YF, 2YE, 2YF/2SA, 2YE/2SD were verified by immunoblot. (**C**) *In vitro* kinase assay using Src and PXN WT, 2SA, or 2SD mutants. (**D**-**G**) Pull-down assay using recombinant Src and PXN mutants. The relative band intensities of PXN were determined by densitometry (**E**) and (**G**). Data are shown as mean ± SEM from 3 independent experiments. (**H**-**K**) Pull-down assays using purified Vinculin or Kindlin-2 and the PXN mutants. ns, not significant; **p < 0.01; ***p < 0.001 by one-way ANOVA.

